# Disruption of c-di-GMP signaling networks unlocks cryptic expression of secondary metabolites during biofilm growth in *Burkholderia pseudomallei*

**DOI:** 10.1101/2021.12.12.472329

**Authors:** Grace I. Borlee, Mihnea R. Mangalea, Kevin H. Martin, Brooke A. Plumley, Samuel J. Golon, Bradley R. Borlee

**Affiliations:** Department of Microbiology, Immunology, and Pathology, Colorado State University, Fort Collins, CO 80523

**Keywords:** diguanylate cyclase, c-di-GMP, biofilm, syrbactin, malleipeptin, *Burkholderia pseudomallei*

## Abstract

The regulation and production of secondary metabolites during biofilm growth of *Burkholderia* spp. is not well understood. To learn more about the crucial role and regulatory control of cryptic molecules produced during biofilm growth, we disrupted c-di-GMP signaling in *Burkholderia pseudomallei,* a soil-borne bacterial saprophyte and the etiologic agent of melioidosis. Our approach to these studies combined transcriptional profiling with genetic deletions that targeted key c-di-GMP regulatory components to characterize responses to changes in temperature. Mutational analyses and conditional expression studies of c-di-GMP genes demonstrates their contribution to phenotypes such as biofilm formation, colony morphology, motility, and expression of secondary metabolite biosynthesis when grown as a biofilm at different temperatures. RNA-seq analysis was performed at varying temperatures in a ΔII2523 mutant background that is responsive to temperature alterations resulting in hypo- and hyper-biofilm forming phenotypes. Differential regulation of genes was observed for polysaccharide biosynthesis, secretion systems, and nonribosomal peptide and polyketide synthase (NRPS/PKS) clusters in response to temperature changes. Deletion mutations of biosynthetic gene clusters (BGCs) clusters 2, 11, 14 (syrbactin), and 15 (malleipeptin) in wild-type and ΔII2523 backgrounds also reveals the contribution of these BGCs to biofilm formation and colony morphology in addition to inhibition of *Bacillus subtilis* and *Rhizoctonia solani*. Our findings suggest that II2523 impacts the regulation of genes that contribute to biofilm formation and competition.

Characterization of cryptic BGCs under differing environmental conditions will allow for a better understanding of the role of secondary metabolites in the context of biofilm formation and microbe-microbe interactions.

**Importance:** *Burkholderia pseudomallei* is a saprophytic bacterium residing in the environment that switches to a pathogenic lifestyle during infection of a wide range of hosts. The environmental cues that serve as the stimulus to trigger this change are largely unknown. However, it is well established that the cellular level of c-di-GMP, a secondary signal messenger, controls the switch from growth as planktonic cells to growth as a biofilm. Disrupting the signaling mediated by c-di-GMP allows for a better understanding of the regulation and the contribution of the surface associated and secreted molecules that contribute to the various lifestyles of this organism. The genome of *B. pseudomallei* also encodes cryptic biosynthetic gene clusters predicted to encode small molecules that potentially contribute to growth as a biofilm, adaptation, and interactions with other organisms. A better understanding of the regulation of these molecules is crucial to understanding how this versatile pathogen alters its lifestyle.

## Introduction

B. pseudomallei is a saprophytic bacterium that switches to a pathogenic lifestyle in a range of hosts causing melioidosis, an often-fatal disease that is common in Southeast Asia, Northern Australia, and other parts of the world (1, 2). Recent published estimates predict approximately 165,000 human cases of melioidosis annually with greater than 50% mortality in 79 countries where the pathogen is endemic (3). The environmental cues that serve as the impetus for *B. pseudomallei* to initiate this lifestyle change from saprophyte to pathogen in coordination with signaling cues are unknown; although, rain, humidity, and wind are thought be drivers of increased *B. pseudomallei* prevalence (4, 5). While these climatic factors are beginning to be defined in the context of the epidemiology of disease transmission, the cues and signal sensing systems in *B. pseudomallei* that contribute to this process are still largely unknown. Bacteria have evolved to sense and respond to their external environment and as a result have developed sophisticated signaling systems to rapidly adjust to their dynamic environment. One such elegant signaling cascade involves c-di-GMP, which has been shown to be an important secondary messenger in numerous bacterial pathogens (6–9).

To better understand c-di-GMP signaling in pathogenic *Burkholderia* spp. (10), we previously characterized 22 transposon insertional mutants predicted to be involved in *B. pseudomallei* 1026b c-di-GMP signaling (11). Curiously, two adjacent transposon mutants in a phosphodiesterase (PDE), *cdpA* (I2284), and a non-canonical HD-GYP protein, I2285, exhibited a reduction in motility (11). We also observed reduced biofilm formation at 30°C and increased biofilm formation at 37°C in a transposon insertion mutant in II0885, which encodes a predicted protein with diguanylate cyclase and phosphodiesterase (DGC/PDE) activity (11). The most significant phenotype with the greatest dynamic range in biofilm response was observed for a transposon insertional mutant of II2523, a predicted diguanylate cyclase (DGC). This mutant exhibited reduced biofilm formation at 30°C, but paradoxically exhibited enhanced biofilm formation at 37°C (11). To further characterize these phenotypes in this study, we created in-frame deletions of *cdpA*, (I2284, PDE), I2285 (degenerate HD-GYP protein), II0885 (hybrid DGC/PDE), and II2523 (DGC) to better understand how each of these genes contributes to c-di-GMP-regulated phenotypes such as biofilm formation and motility in *B. pseudomallei*. In some cases, we constructed site-directed mutations in these genes to identify specific amino acids that contribute to the phenotypes observed. Additionally, we generated a series of mutant strains in the select agent excluded and attenuated strain Bp82 of *Burkholderia pseudomallei*. Bp82 is a Δ*purM* mutant of *B. pseudomallei* strain 1026b that is deficient in purine biosynthesis and is unable to replicate in human cells and has previously been shown to be fully attenuated in hyper susceptible animal models, which include Syrian hamsters and immune deficient mice (12, 13). The Bp82 deletion strains and their isogenic complements in addition to site-direct mutations in these genes provides a tool kit to safely study how these genes and changes in targeted amino acids contribute to c-di-GMP signaling, biofilm formation, and secondary metabolite production in a BSL2 lab. These strains also afford the opportunity to better understand the physiology of this bacterium regarding how it responds to temperatures that it would encounter growing as a saprophyte and during infection of a human host. By perturbing c-di-GMP signaling we can study the biofilm matrix and surface associated components that are differentially expressed in addition to cryptic metabolites that are not expressed in wild-type bacterial cultures that are grown under standard laboratory conditions.

A secondary goal of this study was to enhance our understanding of c-di-GMP-mediated regulation under biofilm inducing growth conditions. We performed RNA-seq and differential gene expression analysis of ΔII2523 in cells grown statically as biofilms at either 28°C or 37°C. In addition to the genes that contribute to biofilm formation (e.g., polysaccharide biosynthetic gene clusters) this analysis also revealed a suite of genes that were differentially regulated that included several NRPS/PKS biosynthetic gene clusters (BGCs). Some of these clusters have been previously characterized, however our mutational approach resulted in unlocking the expression of BGCs that have been previously described as cryptic with unknown functional roles. Recently, there has been a lot of interest in characterizing the metabolites produced by *Burkholderia* spp. and understanding their roles during growth, competition, survival, and infection of hosts (14–17). The BGC notation used herein was originally described by Biggins et al. (14) and we have further characterized some of these BGCs in this research. To better evaluate the role of these cryptic BGCs, we generated combinatorial mutants of ΔII2523 with BGC cluster 2 (unknown nonribosomal peptide synthetase [NRPS]), cluster 11 (unknown NRPS), cluster 14 (syrbactin), and cluster 15 (malleipeptin) to evaluate the contribution of these BGCs to biofilm formation and production of antimicrobial compounds. Overall, we sought to further delineate the complexity of c-di-GMP signaling and the potential contribution of cryptic secondary metabolism to various c-di-GMP-controlled phenotypes in *B. pseudomallei*.

## Results

### Contribution of *B. pseudomallei* c-di-GMP genes to biofilm formation

We have previously demonstrated that a *B. pseudomallei* 1026b transposon mutant inserted in Bp_1026b II2523 produces significantly less biofilm at 30°C than the wild type which is to be expected with the loss of a diguanylate cyclase; however, at 37°C the II2523 transposon mutant is a hyperbiofilm producer in comparison to the wild type, which is in contradiction to the expected result from a loss of a diguanylate cyclase (11). To provide additional experimental support for this finding, in-frame deletion mutants of ΔII2523 were generated in multiple backgrounds, which recapitulated the temperature responsive biofilm phenotypes in *B. pseudomallei* 1026b (Fig 1A-B) and the isogenic select agent excluded Δ*purM* strain *B. pseudomallei* Bp82 (Fig 10A-B). Temperature-dependent biofilm formation phenotype of the ΔII2523 mutant could be in part attributed to c-di-GMP levels, which are elevated at 37°C and diminished at 30°C in comparison to the wild type (Fig 2). Decreased biofilm phenotype of ΔII2523 at 30°C could be rescued with the Δ*cdpA* or with the Δ*cdpA*-I2285 (I2284-I2285) mutant but not with ΔI2285 alone suggesting that the loss of the *cdpA* phosphodiesterase is sufficient to presumably elevate c-di-GMP levels in the ΔII2523 mutant (Fig 1A). Loss of II0885, which is predicted to contain two MHYT, an EAL, and GGDEF domains and is most closely related to CdpA (38% identity at the amino acid level) was not able to rescue the ΔII2523 biofilm phenotype suggesting that this protein does not work in cooperation either directly or indirectly with II2523 at 30°C (Fig 1A). Interestingly, Δ*cdpA*, ΔI2285, and Δ*cdpA*-I2285 significantly reduced biofilm formation as compared to the wild type in addition to reducing biofilm formation in the ΔII2523 Δ*cdpA*-I2285 hyper biofilm forming background when compared with ΔII2523 at 37°C (Fig 1B). Neither Δ*cdpA* nor ΔI2285 mutants relieved the hyper biofilm of ΔII2523 at 37°C (Fig 1B). Interestingly, ΔII0885 significantly reduced the ΔII2523 hyper biofilm formation phenotype resulting in levels that were more similar to wild-type levels suggesting that II0885 contributes to the hyper biofilm forming phenotype at 37°C (Fig 1B). The deletion of II0885 had no effect in the quadruple mutant, Δ*cdpA*-I2285 ΔII0885 ΔII2523, which was identical to the triple mutant, Δ*cdpA*-I2285 ΔII2523 at 37°C (Fig 1B), however, deletion of II0885 did decrease biofilm formation in all mutant combinations tested at 30°C (Fig 1A).

**Figure 1.**
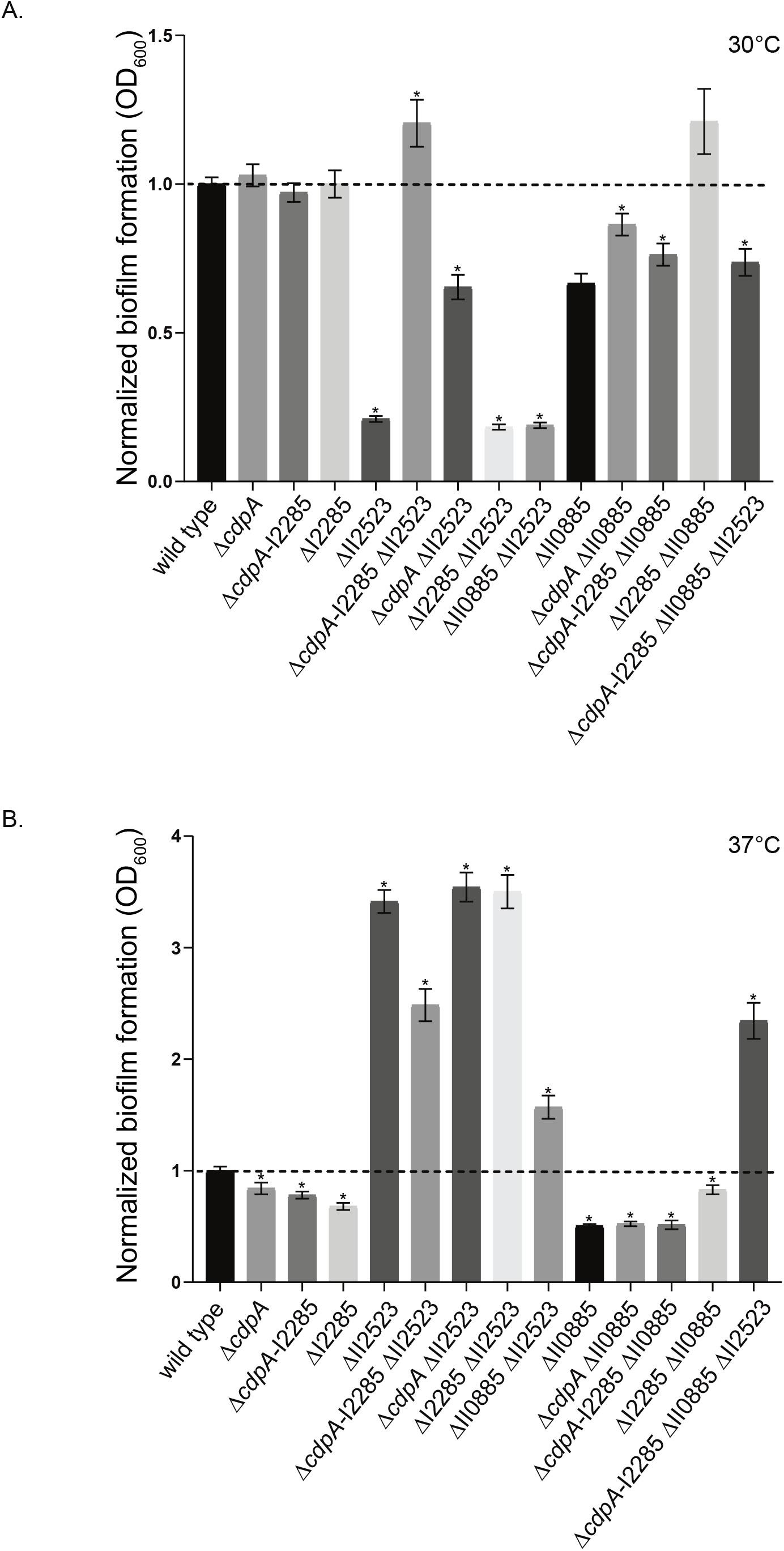
Biofilm formation of *B. pseudomallei* 1026b c-di-GMP deletion mutants. Wild type, single, double, triple, and quadruple mutants were grown statically at 28°C (A) and 37°C (B) for 24h. Data are representative of three independent experiments. Asterisks indicates a significance difference as obtained with a Student’s t-test utilizing Bonferroni correction (p<0.002) to account for multiple comparisons (n=13).

**Figure 2.**
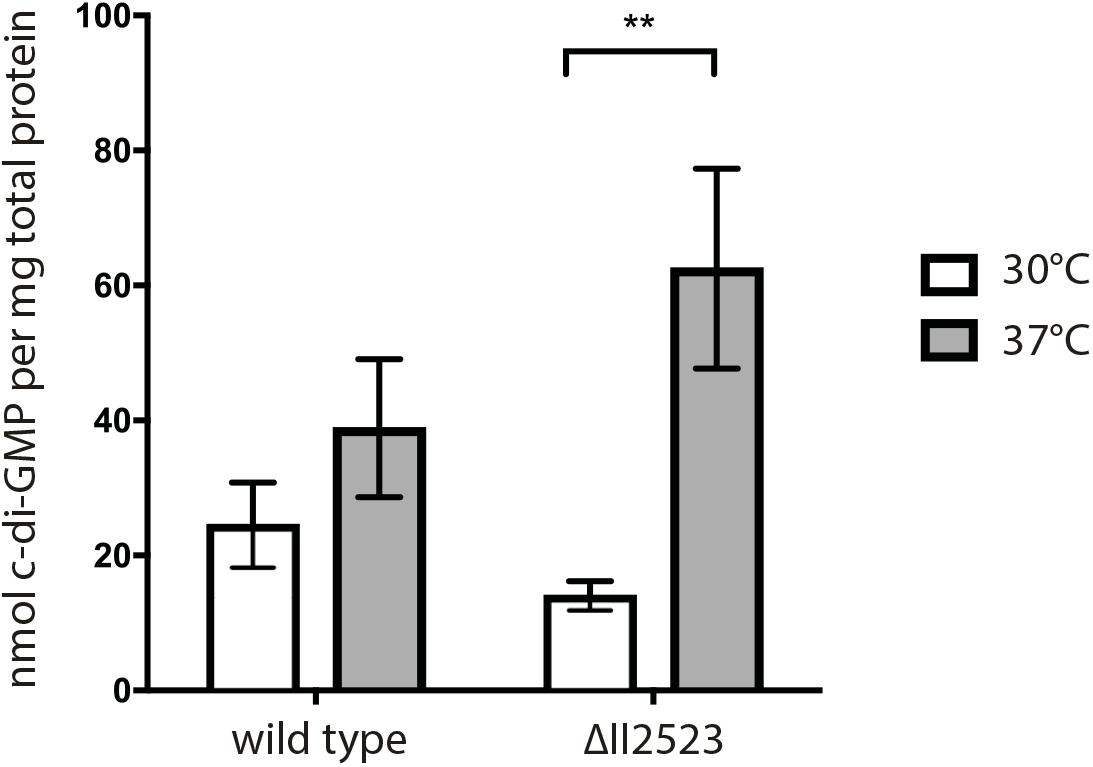
Quantification of c-di-GMP levels of *B. pseudomallei* 1026b and ΔII2523 grown statically at 30°C and 37°C. p<0.01 need info on stats. Statistical significance was determined using the Sidak-Bonferroni method across multiple T-tests (** = p<0.01). Error bars indicate standard error for three technical replicates. C-di-GMP extractions were repeated on separate days using two biological replicate cultures for each strain and temperature condition, with three technical replicates each.

### Contribution of *B. pseudomallei* c-di-GMP genes to motility

Differences in *B. pseudomallei* swimming were observed in plate-based motility assays. Δ*cdpA*, ΔI2285, Δ*cdpA*-I2285, Δ*cdpA* ΔII0885, ΔI2285 ΔII0885, and the triple mutant Δ*cdpA*-I2285 ΔII0885 all exhibited a decrease in swimming motility at both 30°C and 37°C (Fig 3A-B). Decreased motility of strains with mutations in *cdpA* or mutations in orthologs of *cdpA* have been noted in several *Burkholderia* spp. (6, 11, 18, 19). Both ΔII2523 and the orthologous Δ*bcam2836* deletion in *B. cenocepacia* H111 exhibit increased motility (11, 19). Loss of II0885 did not alter the motility phenotypes of either the Δ*cdpA* or ΔI2285 single mutants suggesting that *cdpA* and I2285 are epistatic to II0885 (Fig 3A-B). The loss of both *cdpA* and I2285 was not additive suggesting that these proteins more than likely function in the same pathway (Fig 3A-B). Furthermore, *cdpA* and I2285 are not co-transcribed during planktonic growth suggesting that these genes are regulated independently of each other under the conditions tested (Fig S1). The deletion of *cdpA* or I2285 did not significantly alter hyper-motility in the ΔII2523 mutant background suggesting that other phosphodiesterases or mechanisms also participate in the signaling that controls swimming motility (Fig 3A-B).

**Figure 3.**
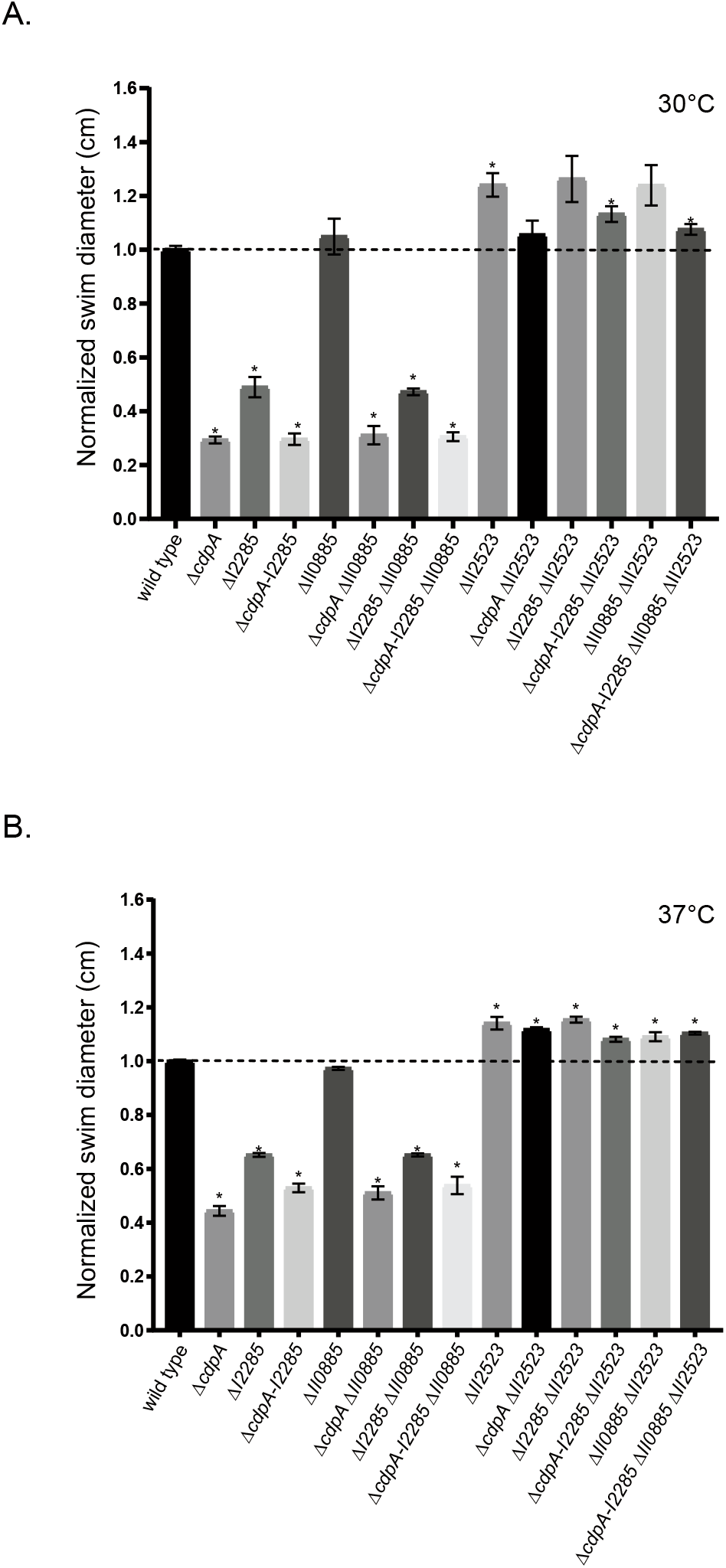
Swimming motility of *B. pseudomallei* c-di-GMP deletion mutants. Swimming motility of the wild type, single, double, triple, and quadruple mutants in 0.3% agar plates. Plates were incubated at 30°C (A) and 37°C (B) for 24h. Asterisks indicates a significance difference as obtained with a Student’s t-test utilizing Bonferroni correction (p<0.002) to account for multiple comparisons (n=13).

### Conditional expression of *cdpA* leads to decreased biofilm formation whereas I2285 results in an increase in biofilm formation in *B. pseudomallei* Bp82

To more rapidly evaluate the contribution of c-di-GMP genes to biofilm formation and the corresponding phenotypes, we constructed our mutants in *B. pseudomallei* Bp82, which is a select-agent excluded avirulent derivative of *B. pseudomallei* 1026b that can be used for research in a BSL2 laboratory (13). Subsequent temperature variation experiments with *B. pseudomallei* Bp82 were conducted at 28°C and 37°C to approximate the temperatures that this opportunistic pathogen would respectively encounter residing in the environment and also during infection of a human host. IPTG-induced expression of *cdpA*, I2285, *cdpA*-I2285, and II0885, but not II2523, resulted in decreased biofilm formation at 28°C suggesting that *cdpA*, I2285, *cdpA-*I2285, and II0885 may function to inhibit biofilm formation at 28°C (Fig 4A). Strikingly, inducible expression of I2285 at 37°C resulted in a significant increase in biofilm formation not observed at 28°C suggesting multiple functions for I2285 that are temperature dependent (Fig 4A-B). Both *cdpA* and *cdpA-*I2285 expression reduced biofilm formation in the wild-type background at 37°C (Fig 4A). Inducible expression of *cdpA* or *cdpA*-I2285 resulted in decreased biofilm formation at 28°C (Fig 4C). Conditional expression of II0885 resulted in decreased biofilm formation at 28°C and 37°C as compared to the wild type while induction of II2523 enhanced biofilm formation as compared to the uninduced control at 37°C (Fig 4B). Conditional expression of either *cdpA*, I2285, or both *cdpA*-I2285, resulted in increased swimming diameter at 28°C and 37°C (Fig 4C and 4D) suggesting that these genes contribute to the regulation of swimming motility (Fig 4C and 4D). Differences in swim motility were more noticeable at 28°C as opposed to 37°C (Fig 4C). This is consistent with *cdpA* encoding for a phosphodiesterase. Conditional expression of II2523, which encodes for a putative diguanylate cyclase, resulted in decreased motility at both temperatures whereas conditional expression of II0885 had no effect (Fig 4C).

**Figure 4.**
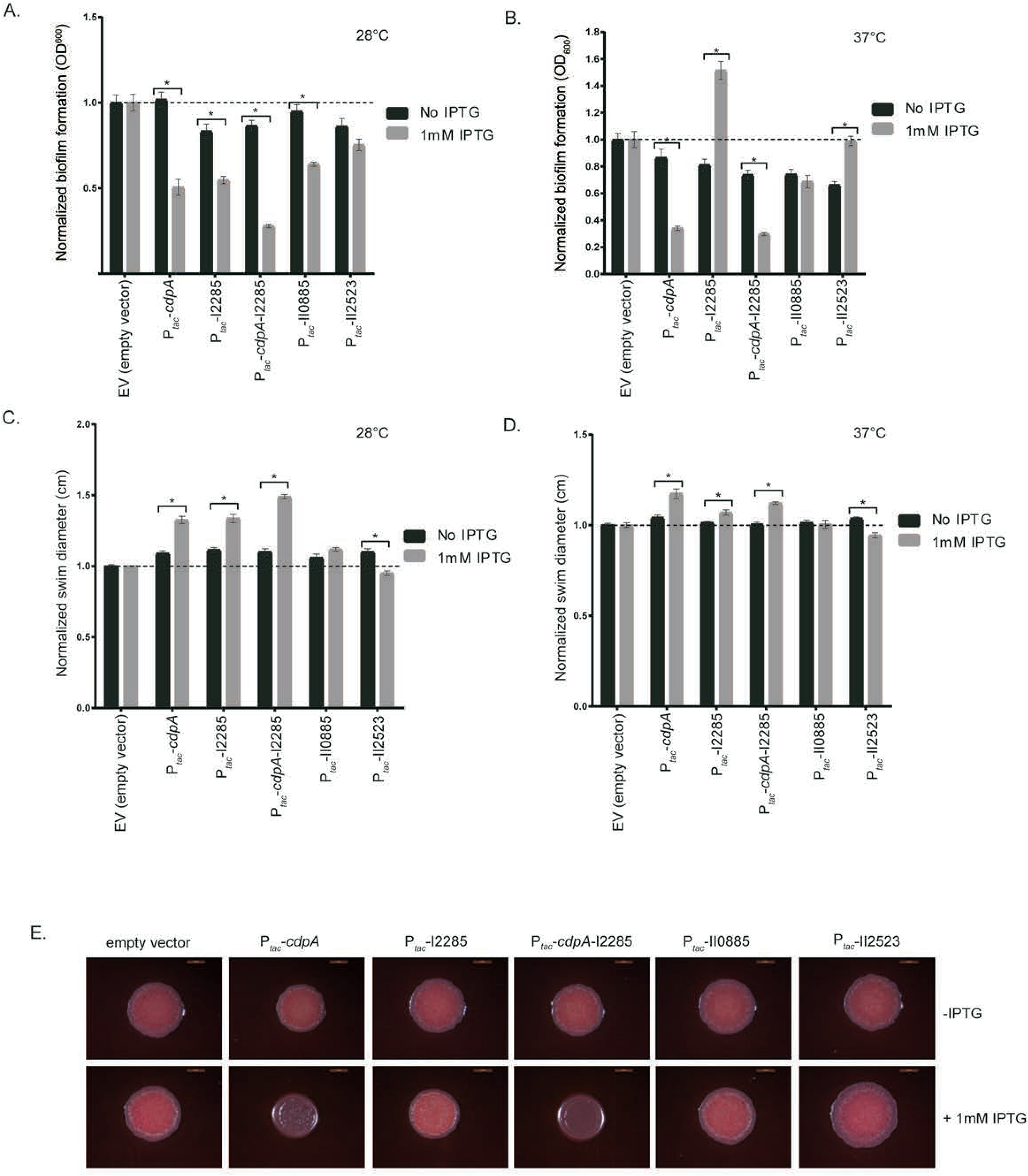
Biofilm formation, swimming motility, and colony morphology of *B. pseudomallei* Bp82 strains conditionally expressing *cdpA*, I2285, *cdpA* I2285, II0885, and II2523. Biofilm assays were incubated at 28°C (A) or 37°C (B) for 24h. Swim assays were incubated at either 28°C (C) or 37°C (D) for 24h. Colony morphology on NAP-A plates incubated at 37°C (E). Images were taken after three days. Conditional expression of c-di-GMP genes was achieved with the addition of 1mM IPTG.

Colony morphology was assessed to characterize phenotypes that are controlled by c-di-GMP (e.g. biofilm-forming capacity, exopolysaccharide production); however, in this study we did not observe striking differences in colony morphology for the c-di-GMP deletion strains grown on either LB, NAP-A or YEM media at either 28°C or 37°C (Fig S2A-F). We also evaluated conditional expression as a means to evaluate the potential function of these genes. *B. pseudomallei* Bp82 strains with inducible expression of *cdpA*, I2285, *cdpA*-I2285, II0885, and II2523 were grown on LB, YEM or NAP-A media with agar and incubated at 28°C and 37°C. There were no discernible differences in colony morphology on LB, YEM at either temperature or NAP-A at 28°C (Fig S4A-C). However, IPTG-inducible expression of *cdpA* or *cdpA*-I2285 resulted in the notable loss of rugosity on NAP-A at 37°C suggesting that *cdpA* can alter colony morphology (Fig 4E).

We also evaluated the activity of these genes using a heterologous approach to evaluate the effect on c-di-GMP signaling mediated phenotypes in *Pseudomonas aeruginosa* PAO1 (20–22). Inducible expression of *cdpA* or *cdpA*-I2285 in either *P. aeruginosa* PAO1 (Fig S3A) or the isogenic hyperbiofilm former PAO1 Δ*wspF* resulted in decreased biofilm formation (Fig S3B). However, motility as measured by swim diameter was not significantly affected by conditional and heterologous expression in *P. aeruginosa* PAO1 (Fig S3C).

### Mapping residues important for activity in CdpA, I2285, and II2523

CdpA (I2284) is a predicted EAL-GGDEF hybrid that retains the canonical EAL domain; but the canonical GGDEF domain is replaced with the ASDFK residues, suggesting that this protein most likely functions solely as a phosphodiesterase rather than as both a diguanylate cyclase and phosphodiesterase (6, 11). A single point mutation in *cdpA* to alter the EAL domain to AAL amino acid motif was constructed to assess the necessity of the EAL domain. Complementation of Δ*cdpA* with the *cdpA*^AAL^ construct did not restore motility to wild-type levels suggesting that the EAL is important for CdpA-mediated motility (Fig 5A). Complementation of Δ*cdpA* with an inducible wild-type *cdpA* decreased biofilm formation suggesting that additional phosphodiesterase activity was responsible for biofilm inhibition (Fig 5B). In addition to the EAL domain, CdpA also contains a PAS domain (11). PAS domains are sensory domains that can perceive signaling cues such as light, FMN, FAD, heme, and others (23). Strikingly, both CdpA and II2523 have PAS domains identified using Pfam analysis (11). CdpA does not retain a highly conserved asparagine but instead has a serine while II2523 retains the conserved asparagine (24). Mutating the serine at position 72 to an alanine in the recombinant CdpA did not alter biofilm and swim motility in the Δ*cdpA* mutant during complementation (Fig 5A-B).

**Figure 5.**
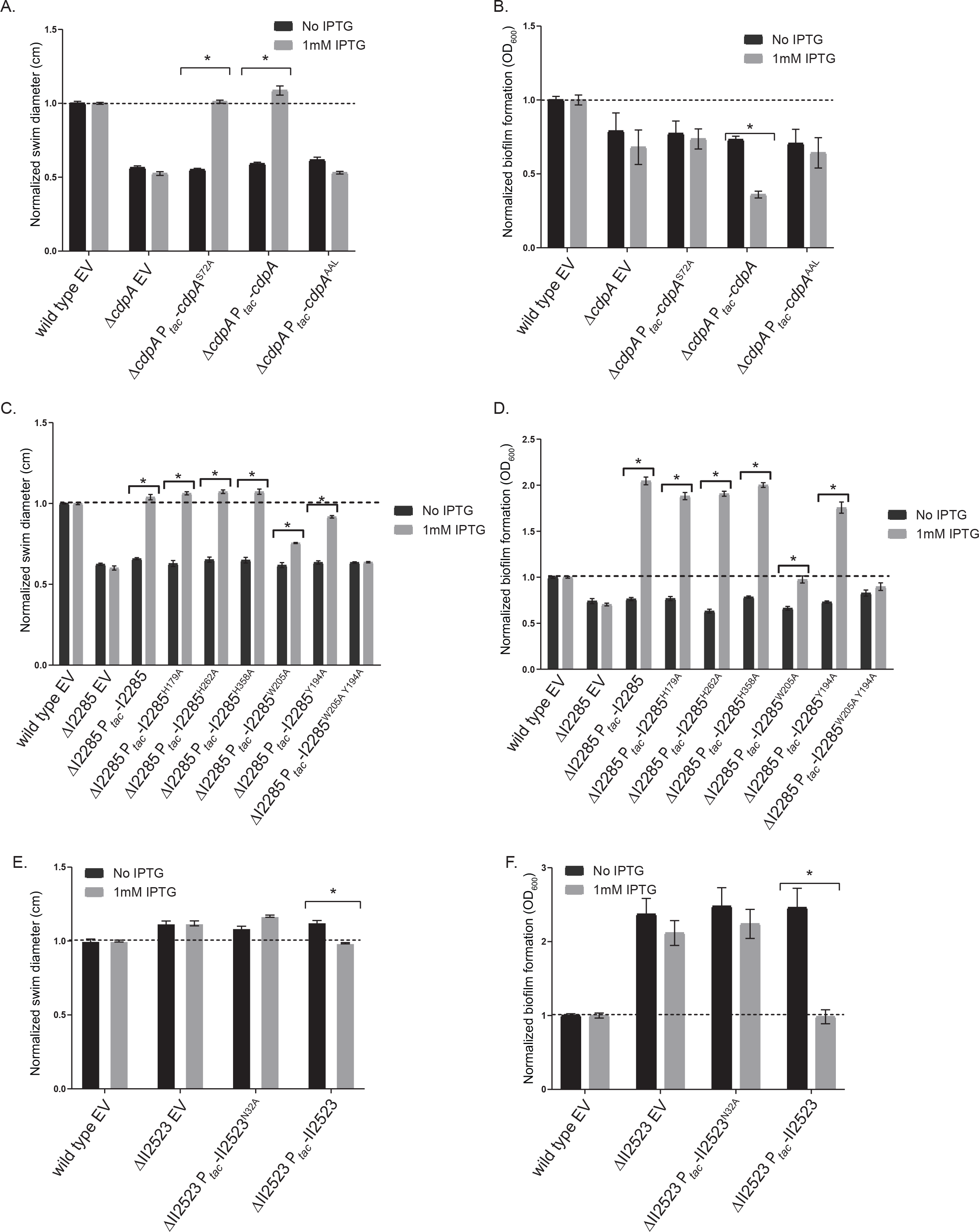
Site-directed mutations in *cdpA*, I2285, and II2523 identify amino acids that are important for swim and biofilm phenotypes in Bp82. Swim (A) and biofilm (B) phenotypes of CdpA PAS4 (S72A) and EAL mutants. Swim (C) and biofilm (D) phenotypes of I2285 mutants. Swim (E) and biofilm (F) phenotypes of II2523 PAS4 (N32A) mutant. 1mM IPTG was used to induce conditional expression. All assays were done at 37°C.

Adjacent to *cdpA* (I2284) lies the gene locus I2285 which encodes for a degenerate HD-GYP protein based on protein alignment; however, we hypothesized that one of the histidine residues, which has been shown to be important in metal binding in HD-GYP proteins, might also be important for I2285 activity (11). We targeted three histidines (H179, H262, and H358) in I2285 for mutagenesis by replacing the respective histidine with alanine. None of these histidines contributed to the swim or biofilm phenotypes of induced I2285 (Fig 5C-D). Sequence alignment of I2285 with GsmR, an HDOD protein from *X. campestris*, revealed additional amino acid residues to target for mutagenesis (Fig S5). Proteins with the HD-related output domain (HDOD) are widespread and can coordinate the signaling that controls chemotaxis (25). Liu et al. have previously shown that mutating a conserved tryptophan in GsmR resulted in loss of complementation in a swim assay (26). Mutating both the highly conserved tryptophan (W205) and a nearby tyrosine (Y194) in I2285 to alanine partially disrupted the increased motility of recombinant I2285 expression in the ΔI2285 mutant (Fig 5C). However, incorporating both mutations (Y194A and W205A) completely disrupted complementation in both swim and biofilm assays (Fig 5C-D). Interestingly, the single mutation of W205A did partially reduce the hyper-biofilm formation of overexpression of recombinant I2285, whereas Y194A did not (Fig 5D). Mutation of the asparagine (position 32) of the PAS domain (11) resulted in a minor increase in ΔII2523 swimming motility although it was not significant (Fig 5E). This same PAS mutation in II2523 did not complement the ΔII2523 biofilm phenotypes at 37°C back to wild-type levels indicating the necessity of this asparagine, whereas full length II2523 was able to fully complement the hyperbiofilm phenotype of II2523 at 37°C (Fig 5F).

### Transcriptional analysis of ΔII2523 and wild-type biofilms at 28°C and 37°C reveals a multitude of differential expression genes among pairwise comparisons

Differential expression analysis of wild-type *B. pseudomallei* Bp82 biofilm formation at 37°C as compared to 28°C exhibited numerous transcript changes after 24 h of static growth (Table S2). We used the DESeq package (27) for RNA-seq differential expression analyses at thresholds of log2 fold change < -1 or > 1 and adjusted p-value < 0.01. At these thresholds, 123 genes were significantly up-regulated with the highest being II1640 (T3SS-3 secretion system) at 44-fold and 318 genes that were significantly down-regulated with the lowest being II1733 (T3SS-2 secretion system) at -123-fold when comparing wild-type biofilm formation at 37°C versus 28°C (Fig 6A, Table S2).

**Figure 6.**
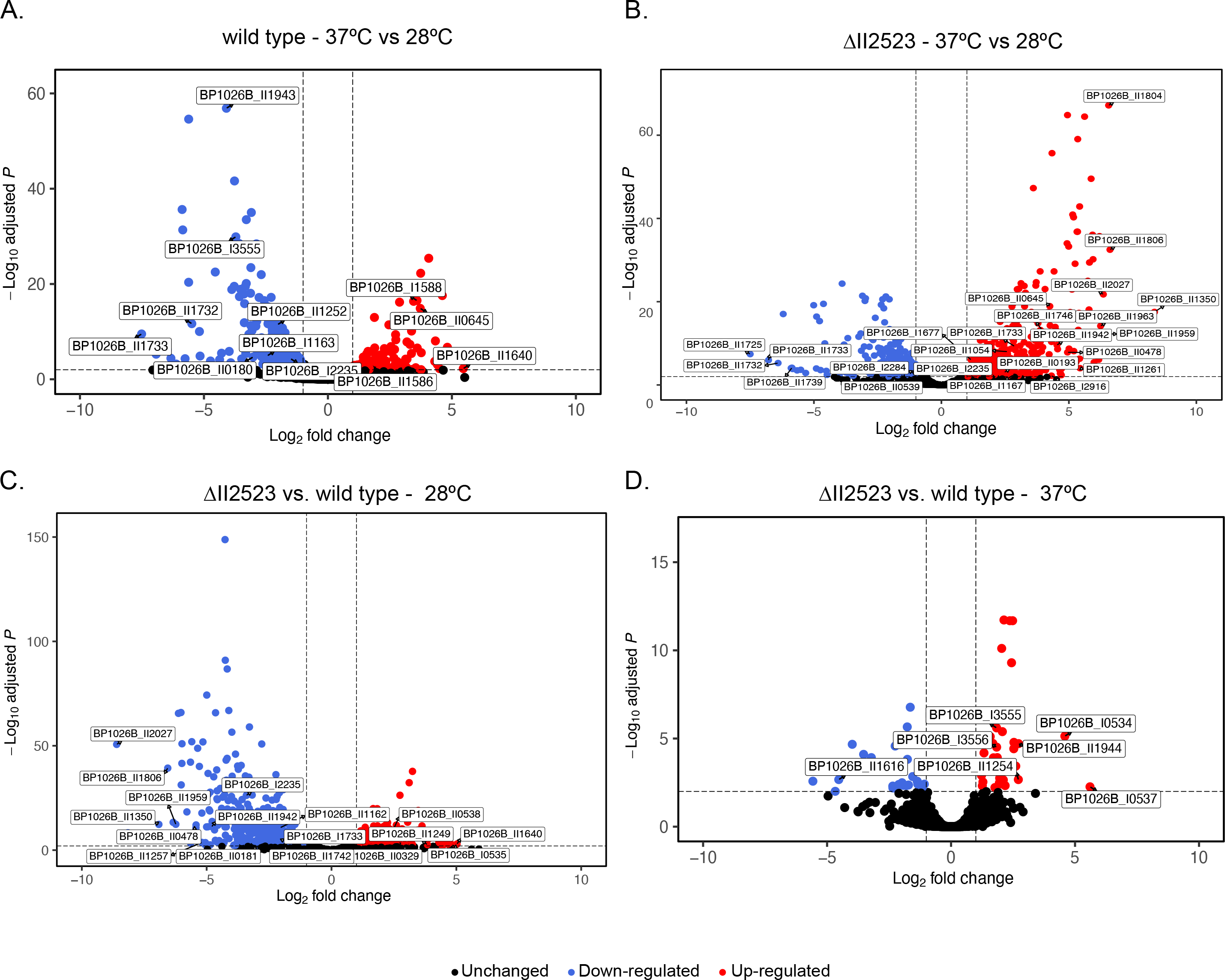
Volcano plots of genes differentially regulated in Bp82 wild type (A) and Bp82 ΔII2523 (B) cells statically grown at 37°C vs. 28°C. Volcano plots of ΔII2523 at 37°C vs. wild type at 37°C (C) and ΔII2523 at 28°C vs. wild type at 28°C (D). Dash lines represent cutoffs for log2 Fold Change < -1 or > 1 (vertical) and adjusted *p-*value significance < 0.01 (horizontal).

Up-regulated genes included pyochelin (II0645, cluster 10), malleipeptin (II1746, cluster 15), malleiobactin (I1735, cluster 1), *bec* biofilm cluster (I2923), T3SS-3, T6SS-3 and T6SS-6 secretion system (Fig 6A and Table S2). Genes down-regulated included ones involved in motility (I3555), T3SS-2, clusters 3 (I1164, unknown), 7 (II0180, isonitrile), 12 (II1252, bactobolin) and 16 (II1943, unknown) NRPS/PKS biosynthesis and a diguanylate cyclase I2235 (Fig 6A and Table S2).

The greatest dynamic range in biofilm formation was observed in the ΔII2523 mutant when grown at temperatures that approximate human body temperature (37°C) as compared to temperatures *B. pseudomallei* encounters growing as saprophyte (28°C or 30°C) (Fig. 1A-B, Fig 10 A-B). Differential expression analyses of all *B. pseudomallei* transcripts indicated there were 443 genes that were up-regulated with the highest being II1350 (syrbactin) at 328-fold and 312 genes that were down-regulated with the lowest being II1725 (T3SS-2) at -181-fold in this comparison (Table S2). Genes that were up-regulated at 37°C as compared to 28°C included those involved in biosynthesis of NRPS clusters 1 (I1733, malleiobactin, siderophore), 2 (I1677, unknown), 3 (I1167, unknown), 10 (II0645, pyochelin), 13 (II1261, unknown), 14 (II1350, syrbactin), 15 (II1746, malleipeptin), 16 (II1942, unknown), *bec* biofilm exopolysaccharide cluster (I2916), T6SS-4, T3SS-3, capsules II (II0478) and III (II1959), *bcaA* (autotransporter), and I2235, a diguanylate cyclase (Fig 6B and Table S2). Genes that were down-regulated included motility genes, 2-alkyl-4-quinolone (II0539, cluster 9) T3SS-2, T3SS-3, and T6SS-5 secretion system, and *cdpA* (I2284) (Fig 6B and Table S2).

At 28°C, ΔII2523 forms considerably less biofilm than the wild type (Fig 1A) (11), a phenotype that would be predicted if II2523 functioned as a diguanylate cyclase. Interestingly, a considerable number of genes (239 significantly up- and 547 down-regulated) were altered when comparing the ΔII2523 to the wild type grown at 28°C (Table S2). Genes that were up-regulated in ΔII2523 as compared to the wild type included capsule IV (I0535), genes involved in motility, malleilactone (II0329, cluster 8), 2-alkyl-4-quinolone (II0538, cluster 9), bactobolin (II1249, cluster 12), T6SS-5, T3SS-3, N-acylhomoserine lactone synthase (*bpsI2*) with the most up-regulated gene being II1640 (part of the T3SS-3 secretion system) (Fig 6C and Table S2). Genes that were down-regulated included: cluster 1 (I1733, malleobactin), cluster 3 (I1162), cluster 7 (II0181, isonitrile), cluster 13 (II1257), cluster 14 (II1350, syrbactin), cluster 15 (II1742, malleipeptin), cluster 16 (II1942), T6SS-4, capsule II (II0478) and III (II1959, *bce-I*), *bce-II* (II1896), and I2235 (diguanylate cyclase) with the most down-regulated gene being II2027 (ortho-halobenzoate 1,2-dioxygenase alpha-ISP protein, OhbB) (Fig 6C and Table S2). A number of these genes in this data set have been previously shown to be differentially regulated by quorum-sensing including cluster 3, 8, 9, 12, 13 along with capsule III (*bce-I*) and *bce-II* (28).

Thirty-six genes were significantly up-regulated when comparing ΔII2523 grown at 37°C to the wild type at 37°C (Table S2). These genes included capsule IV, bactobolin biosynthesis (cluster 12), cluster 16 (unknown NRPS) and motility, with the most up-regulated gene being I0537 (part of capsule IV) at 48.6-fold (Fig 6D). Twenty-six genes, mainly T3SS-3, were down-regulated when comparing ΔII2523 vs wild-type biofilms at 37°C with one of the most down-regulated genes being II1616 (a component of the T3SS-3 secretion system) at -23.1-fold (Fig 6D and Table S2). Although considerably fewer transcripts were differentially expressed when comparing ΔII2523 grown at 37°C to the wild type at 37°C, it is evident that this gene likely contributes to key physiological aspects of *B. pseudomallei* at this host-associated temperature.

### Visualization and validation of temperature-dependent global expression trends reveals significant differentiation in key *B. pseudomallei* functional clusters

Using the Webserver for Position Related data analysis of gene Expression in Prokaryotes (WoPPER) (29) to visualize fold change data output from DESeq2, we identified differential regulation of NRPS clusters and biofilm-associated exopolysaccharide clusters across both *B. pseudomallei* chromosomes (Fig 7). Wild-type cells grown as biofilms at 37°C compared to 28°C exhibited differential expression of motility-associated clusters, a pili cluster, the *Burkholderia* exopolysaccharide cluster (bec), NRPS clusters 1 (malleobactin) and 3, and T6SS-6 on chromosome I, while NRPS clusters 7 (isonitrile), 10 (pyochelin), 15 (malleipeptin), and 16, along with T3SS-2, T3SS-3, and T6SS-3 were differentially affected on chromosome II (Fig 7A). A comparison of ΔII2523 at 37°C vs. 28°C revealed some overlap of affected clusters on both chromosomes, yet clusters 13 and 14 (syrbactin) as well as the biofilm-associated clusters CPSIII (*bce-*I), *bce-*II, and CPSII represented features upregulated specifically in the ΔII2523 background (Fig 7B). Further pairwise comparisons of ΔII2523 vs. the wild type at 37°C (Fig 7C) and ΔII2523 vs. the wild type at 28°C (Fig 7D) showed multiple NRPS, biofilm-associated, and capsule clusters differentially expressed especially on chromosome II, a trend we previously observed in *B. pseudomallei* supplemented with extracellular N-oxide signaling molecules (30). The graphical WoPPER analyses provide visual validation of RNA-seq dataset pairwise comparisons as well as additional orientation for the DESeq2 differential expression output (30).

**Figure 7.**
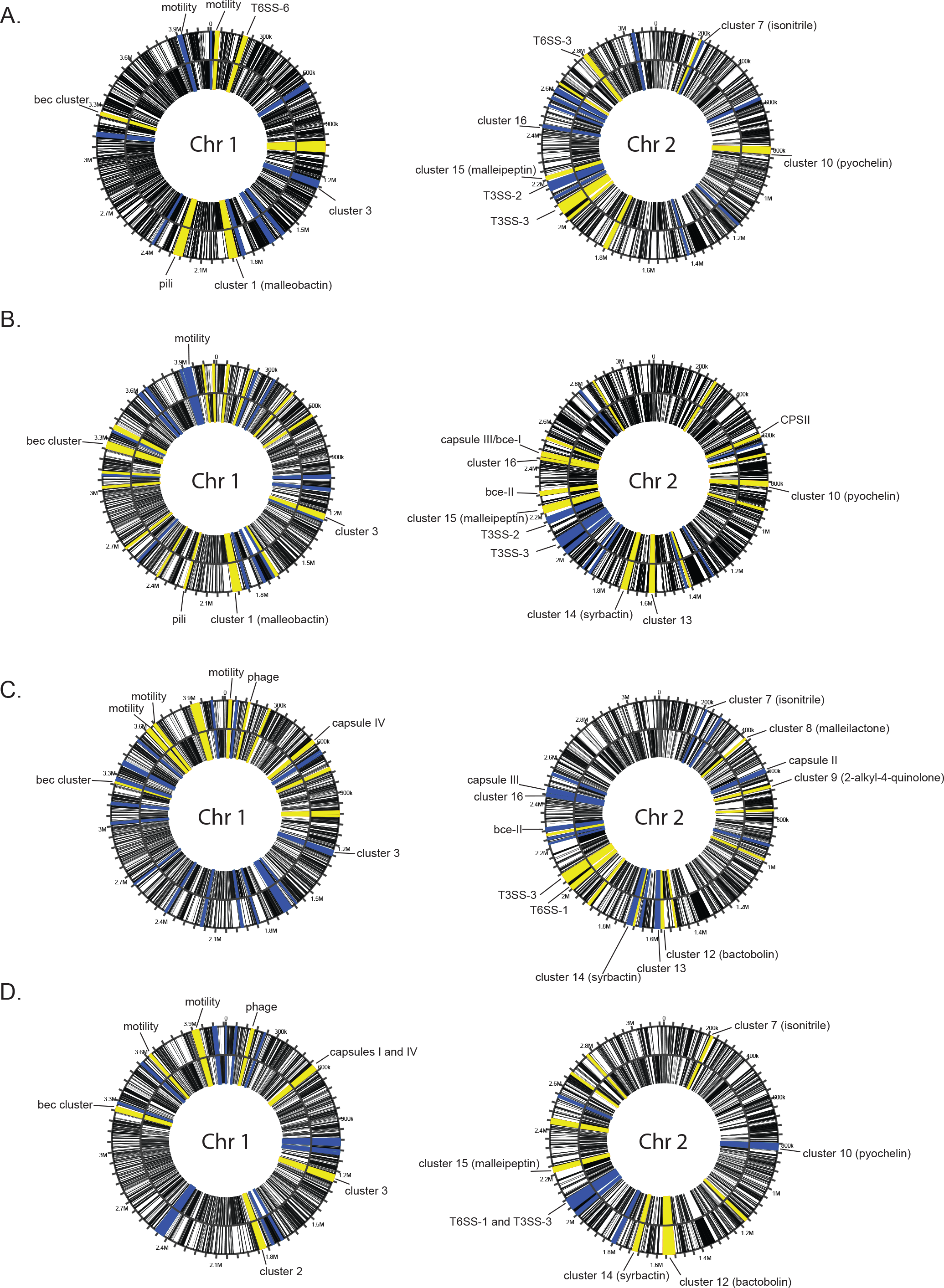
WoPPER analysis of gene clusters differentially regulated in Bp82 wild type and Bp82 ΔII2523 by chromosome. (A) Comparisons of wild-type cells at 37°C vs. 28°C, (B) ΔII2523 at 37°C vs. 28°C, (C) ΔII2523 at 37°C vs. wild type at 37°C, and (D) ΔII2523 at 28°C vs. wild type at 28°C. Yellow indicates gene clusters that were up-regulated and blue indicates gene clusters that were down-regulated.

Greater visual detail for these datasets and pairwise comparisons was achieved via heatmap analysis of complete predicted NRPS clusters (1-3, 5, and 7-16) and biofilm-associated clusters (CPSI, CPSII, CPSIII/*bce-*I, CPSIV, *bce-*II, and *becA-R*) (Fig S6). For the heatmap analyses, raw fold change values from DESeq2 were used as input for all transcripts spanning the predicted clusters (BGC cluster table), except for cluster 4, which is not present in *B. pseudomallei* 1026b, and cluster 6, which is only one gene. The most striking patterns of differential regulation for complete clusters were evident for cluster 13, cluster 14 (Syrbactin), CPSII, CPSIII/*bce-*I, and *bce-*II, where ΔII2523 is similarly implicated in large negative shifts of expression especially at 28°C (Fig S6). Cluster 14 (syrbactin) was most strongly affected, at an average 82.4-fold up-regulated for the entire cluster when comparing ΔII2523 at 37°C vs. ΔII2523 at 28°C and -33.8-fold down-regulated when comparing ΔII2523 at 28°C vs. the wild type at 28°C (Fig S6). Similarly strong trends were observed for the entire clusters CPSIII/*bce*-I and *bce-*II, which were up-regulated at average fold changes of 47.2 and 35.9, respectively, when comparing ΔII2523 at 37°C vs. ΔII2523 at 28°C, and down-regulated at average fold changes of -43.2 and -22.1, respectively, when comparing ΔII2523 at 28°C vs. the wild type at 28°C (Fig S6).

Representative transcripts from gene clusters were further validated via quantitative real-time PCR, which revealed similar expression trends when compared to fold-change differences between sample groups (Fig 8). Although differential abundances calculated as a ratio of logs in pairwise comparisons and quantitative real-time PCR provides a value of one condition to the normal level, these complementary methods serve as further confirmation of RNA-seq analyses. Transcript levels relating to CPSII (II0478) and CPSIII (*bce-I*, II1965) were relatively reduced in ΔII2523 at 28°C, 0.029 and 0.010, respectively (Fig 8). The capsule-associated loci, II0478 and II1965, were significantly down-regulated in a similar fashion when comparing ΔII2523 to the wild type at 28°C; -43-fold and -39-fold change reduction, respectively. II1799, part of the *bce-II* gene cluster, was decreased in ΔII2523 at 28°C at a relative transcript level of 0.025 as compared to a relative transcript level of 2 for ΔII2523 grown at 37°C (Fig 8). RNA-seq evaluation of II1799 revealed a -23.9-fold reduction in expression when comparing ΔII2523 to the wild type at 28°C and a modest increase of 1.2-fold when comparing ΔII2523 to the wild type at 37°C. *fliC* (I3555) expression for the wild type grown at 37°C (Fig 8) was reduced 0.04 relative to the wild type, which agrees with the RNA-seq dataset that indicated a -13-fold change for *fliC* at 37°C compared to 28°C. Capsule IV (I0530) expression was elevated in the ΔII2523 mutant grown at 28°C and less so at 37°C (Fig 8). Correspondingly, I0530 was up-regulated 18-fold and 4-fold when comparing ΔII2523 to the wild type at 28°C and 37°C, respectively, in the RNA-seq data sets (Fig 8). Transcript levels for the EAL/GGDEF hybrid protein-encoding locus, I2928, adjacent to the *bec* biofilm associated gene cluster was down-regulated in the wild type grown at 37°C (0.022 relative transcript) and ΔII2523 grown at 28°C (0.023 relative transcript) (Fig 8). Relative transcript abundances for another c-di-GMP gene, I2284 (*cdpA*), was up 2.6-fold in the ΔII2523 mutant grown at 28°C. Thus, the relative transcript abundances for loci associated with biofilm formation follow a similar trend to the fold change expression differences observed in the comparative RNA-seq datasets across multiple conditions.

**Figure 8.**
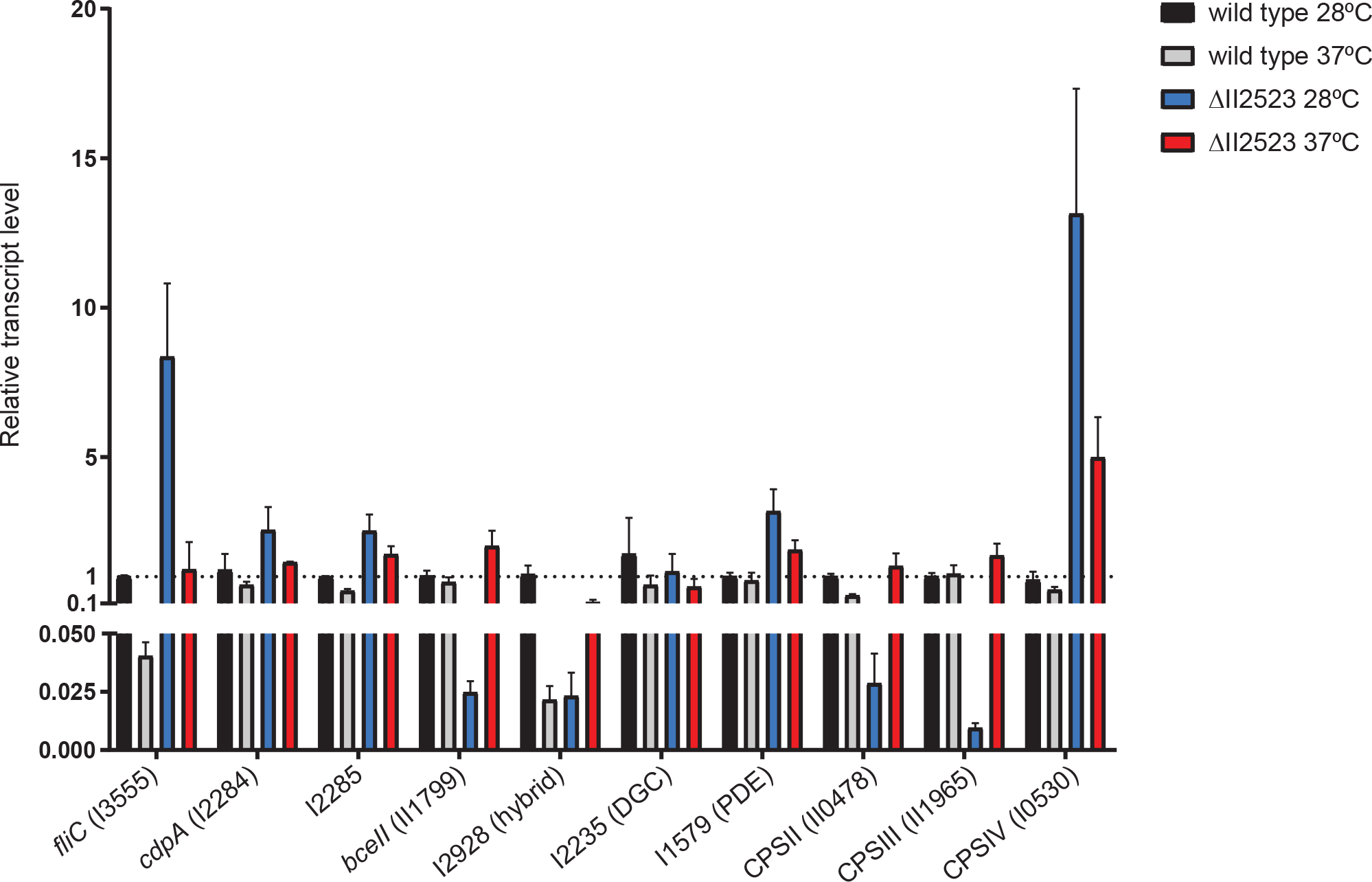
Gene expression (qRT-PCR) of *fliC* (I3555), *cdpA* (I2284), I2285, *bce-II*, (II1799), I2928 (hybrid), I2235 (DGC), I1579 (PDE), capsule II (II0478), capsule III (II1965), and capsule IV (I0530) of biofilm cells from ΔII2523 vs the wild type grown at either 28°C or 37°C. RNA samples for qRT-PCR assays were isolated after 24 h in identical conditions as for the RNA-seq data sets.

### Differential expression of diguanylate cyclase I2235 at varying temperatures reflects a network of c-di-GMP regulation by II2523

Four genes previously predicted to be involved in c-di-GMP signaling in *B. pseudomallei* (11) were differentially expressed passing significance thresholds in our analyses (log2FoldChange < -1 or > 1 and adjusted p-value < 0.01) (Table S2). The phosphodiesterase *cdpA* was down-regulated -2.2 fold in conditions of elevated c-di-GMP production (ΔII2523 at 37°C) compared to low c-di-GMP conditions (ΔII2523 at 28°C). I2235, a predicted diguanylate cyclase, was up-regulated 3.8-fold in the same pairwise comparison (ΔII2523 at 37°C vs ΔII2523 at 28°C) (Table S2). In comparing wild-type Bp82 grown at 37°C vs. 28°C, only the diguanylate cyclase I2235 was down-regulated -2.8-fold (Table S2). This diguanylate cyclase, I2235, was also down-regulated -10.6-fold when comparing ΔII2523 vs. the wild type grown at 28°C while both *cdpA* (I2284) and the adjacent HD-like gene, I2285, as well as I3233 which encodes for an ortholog of the flagellar brake protein, YcgR were both up-regulated 3.6-fold and 2.6- fold, respectively (Table S2). Interestingly, none of the c-di-GMP genes were differentially expressed at our significance threshold when comparing ΔII2523 vs. the wild type at 37°C (Table S2).

### Deletions in BGCs alter colony morphology and biofilm formation in the wild type and ΔII2523

RNA-seq revealed differential regulation of numerous BGCs suggesting that many of these may only be produced during growth as a biofilm and could also be regulated by c-di-GMP (see supplemental table 3 for a description of BGCs). Four of these BGCs (Clusters 2, 11, 14, and 15) have been characterized to varying degrees in *B. pseudomallei* (14). Cluster 2 (I1663-1681) is predicted to encode for a five amino acid lipopeptide that has yet to be characterized in detail (14), but was upregulated in only ΔII2523 at 37°C vs. 28°C and vs. the wild type at 37°C (Fig S6). Cluster 11 (II1089-1108) encodes for an unknown and predicted NRPS/PKS BGC, which was down-regulated in both the wild type and ΔII2523 at 37°C vs. 28°C (Fig S6). Cluster 14 (II1345-1353) encodes the production of syrbactin, a known eukaryotic proteasome inhibitor, which was down-regulated when comparing ΔII2523 at 28°C vs. the wild type at 28°C but was up-regulated in ΔII2523 at 37°C vs. ΔII2523 at 28°C as well as in the ΔII2523 at 37°C vs. the wild type at 37°C conditions (Fig S6). Cluster 15 (II1742-II1746) encodes for malleipeptin, a lipopeptide and a potential biosurfactant (14). Genes for malleipeptin biosynthesis were up-regulated for the wild type and ΔII2523 at 37°C vs. 28°C but down regulated when comparing ΔII2523 and the wild type at 28°C (Fig S6). Together, these data suggest that BGCs contribute to *B. pseudomallei* biofilm formation; however, the exact mechanism of how these secondary metabolites contribute to biofilm formation has yet to be elucidated. We generated a series of deletion mutants of clusters 2, 11, 14, and 15 in both the wild type and ΔII2523 genetic backgrounds to evaluate the contribution of these secondary metabolites to biofilm formation. The loss of cluster 14 (syrbactin) in either the wild type or the ΔII2523 background resulted in smooth colony morphology on NAP-A, YEM, or LB plates regardless of temperature with the notable exception of NAP-A at 37°C (Fig 9A-B and Fig S7). Most of the strains were rugose (wrinkly) on YEM agar at 28°C with the exception of Δcluster 14 mutants (Fig 9A). Cluster 2 deletion mutants were hyper wrinkly at 28°C but the Δcluster 2 in the wild type lost all rugosity at 37°C, while ΔII2523 Δcluster 2 retained some rugosity (Fig 9A). When the media conditions were altered, additional strains (Δcluster 2, ΔII2523 Δcluster 2, Δcluster 11, Δcluster 14, and ΔII2523 Δcluster 14) exhibited smooth colony morphology and increased pigmentation at 28°C as opposed to 37°C on NAP-A (Fig 9B). Interestingly, we noted that ΔII2523 Δcluster 11 reproducibly produced increased radial growth on NAP-A at 37°C compared to other mutants on the same media (Fig 9B).

**Figure 9.**
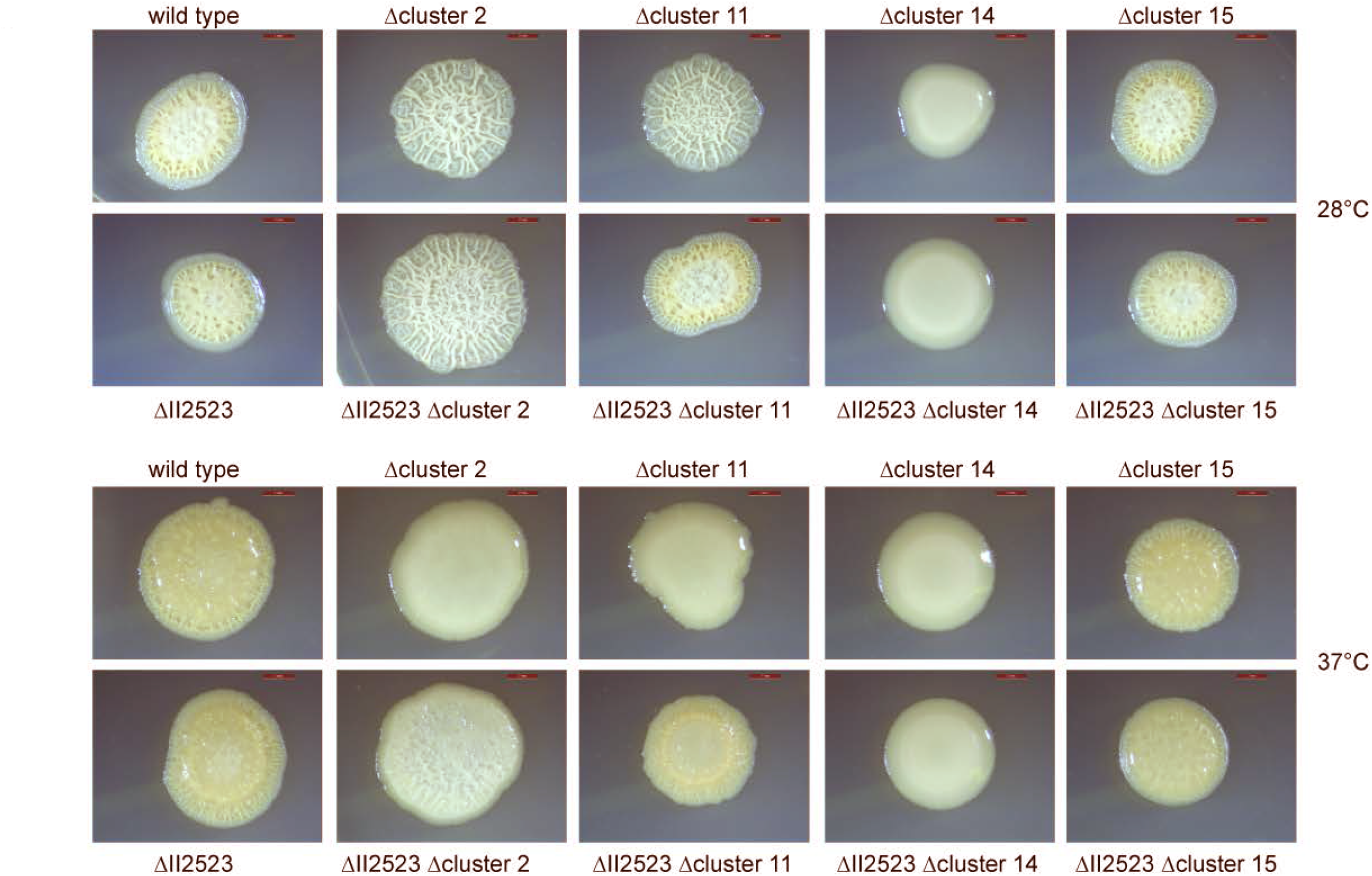
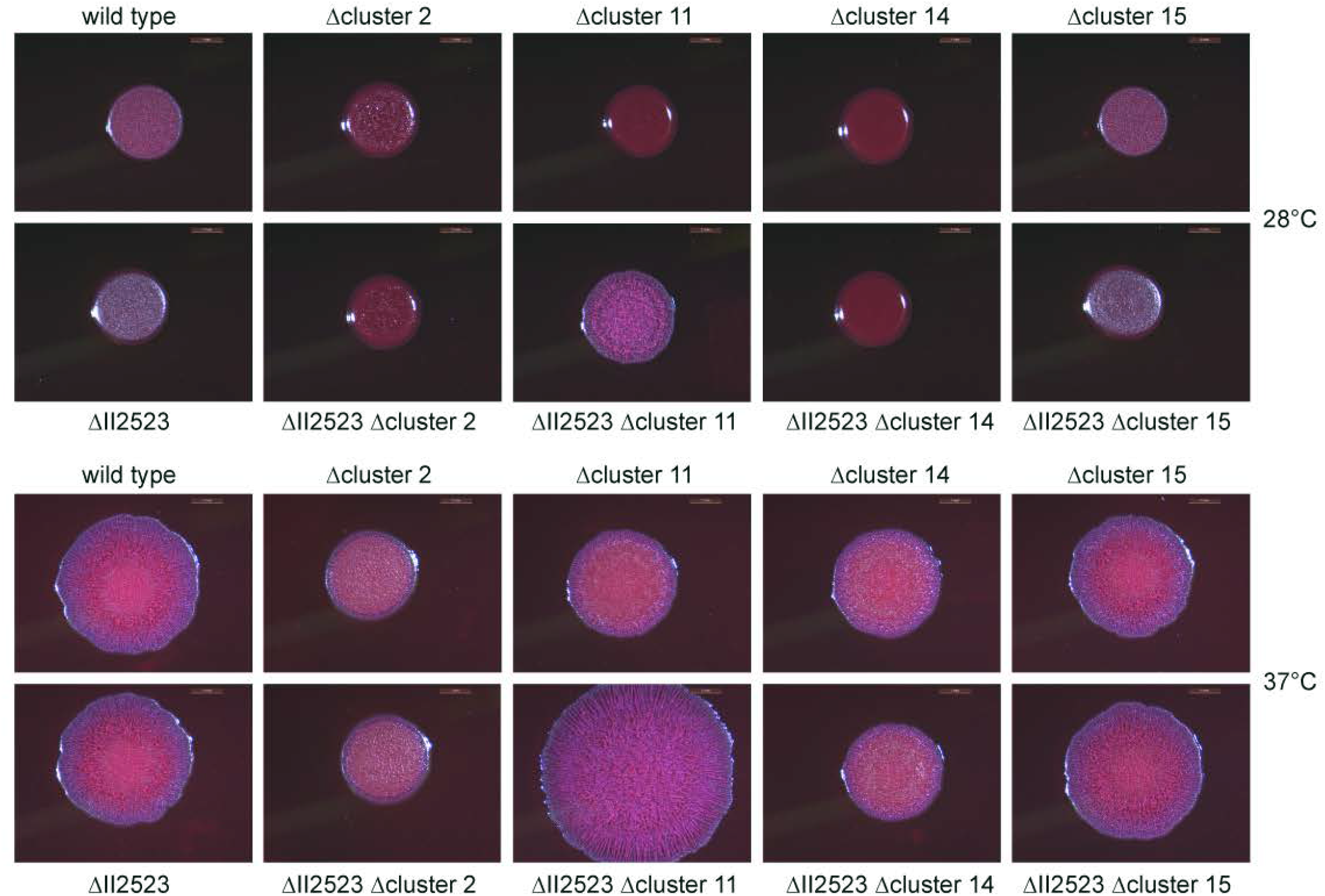
Colony morphology of NRPS/PKS mutants in Bp82 wild type or Bp82 ΔII2523 backgrounds on different media. Spots were grown on YEM (A) or NAP-A (B) at 28°C or 37°C. Images were taken after four days of growth. Scale bar represents 2mm.

**Figure 10.**
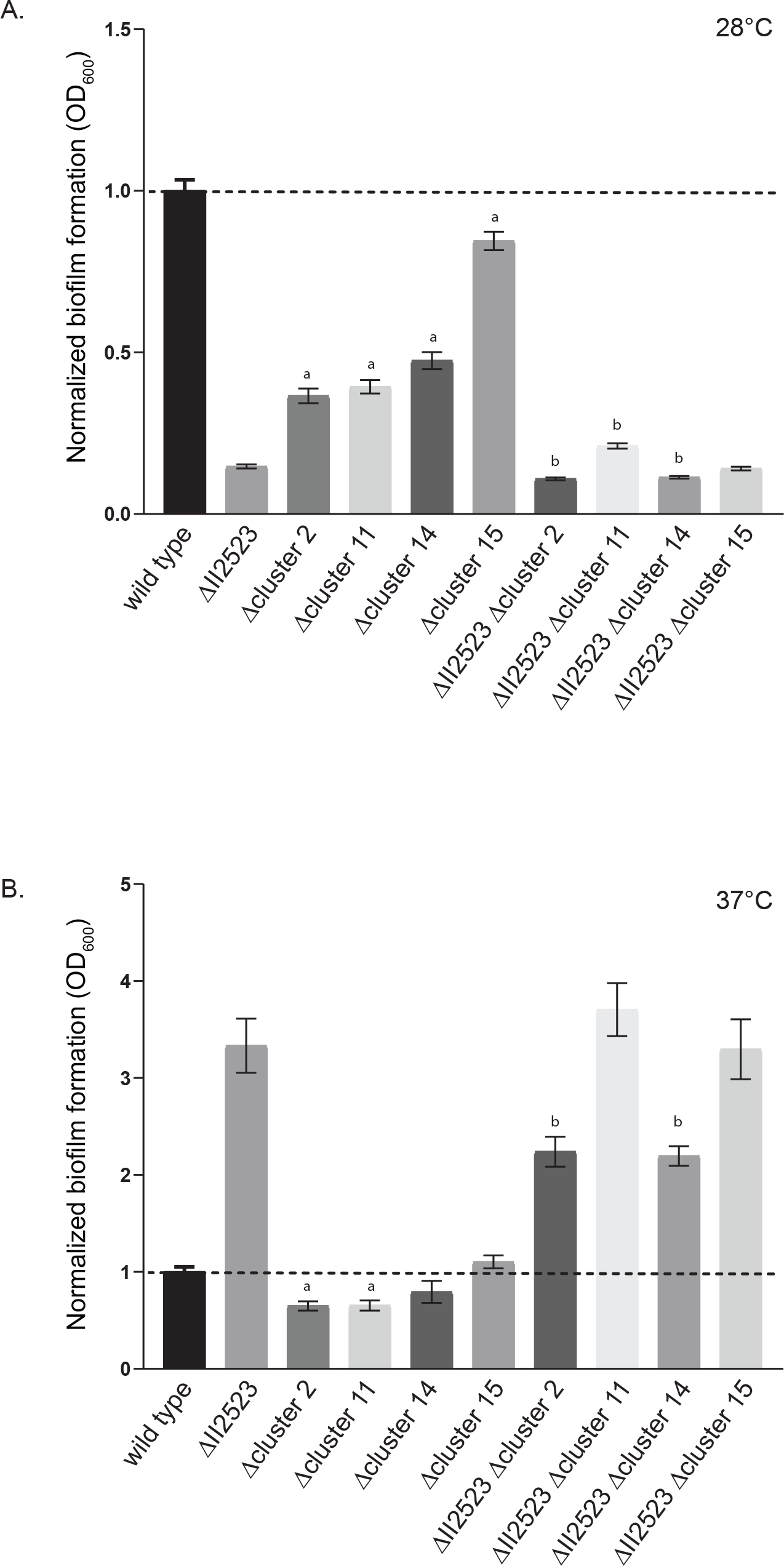
Biofilm formation of BGC Bp82 mutants either 28°C (A) or 37°C (B) for 24h. Lowercase “a” indicates a statistical difference from wild type while a lowercase “b” indicates a statistical difference from ΔII2523.

As noted by Biggins and colleagues, malleipeptin (cluster 15) is a biosurfactant and thus might play a role in biofilm formation (14). Deletion strains of BGCs were evaluated for biofilm formation at both at 28°C and 37°C (Fig 10A-B). Deletion of BGCs 2, 11, 14 (syrbactin), and BGC 15 (malleipeptin) in the wild-type Bp82 background resulted in decreased biofilm formation although to varying degrees at 28°C with the loss of cluster 15 exhibiting the smallest decrease (Fig 10A). Loss of clusters 2 and 14 decreased biofilm formation in the ΔII2523 mutant background while the loss of cluster 11 slightly enhanced biofilm formation in the ΔII2523 mutant at 28°C (Fig 10A).

Only Δcluster 2 and Δcluster 11 exhibited less biofilm formation than wild type at 37°C (Fig 10B). Double mutants ΔII2523 Δcluster 2 and ΔII2523 Δcluster 4 (syrbactin) exhibited decreased biofilm formation at 37°C (Fig 10B) as compared to ΔII2523. Notably, both ΔII2523 ΔBGC2 and ΔII2523 Δcluster 14 exhibited smooth colony morphology depending on the media and temperature (Fig 9A-B and Fig S7).

Appropriately, two NRPS clusters (BGCs) encoding bactobolin (cluster 12) and an unknown metabolite (cluster 16) were up-regulated in ΔII2523 as compared to the wild type at 28°C as observed in the RNA-seq data (Fig 6D andTable S2). Bactobolin from *B. thailandensis* is a quorum-sensing mediated antibiotic effective against *B. subtilis* (31). Bactobolin from *B. pseudomallei* has been reported as a toxic polyketide-peptide hybrid molecule that interferes with host protein synthesis and is sensed by the bacterial feeding nematode, *Caenorhabditis elegans* (32). This potentially indicates that bactobolin could be an important molecule regulated by c-di-GMP that is secreted during biofilm formation to protect against predation and kill competing bacteria.

### BGCs contribute to growth inhibition of *B. subtilis* and *R. solani*

Characterization of the biological activity of the secondary metabolites produced by BGCs 2, 11, 14 (syrbactin), and 15 (malleipeptin) from *B. pseudomallei* has been limited by the cryptic nature of the molecules encoded by these BGCs (14, 16). Deletion of BGCs 2, 11, 14, and 15 were assessed in competition with *Bacillus subtilis* and *Rhizoctonia solani* (a soil-borne fungal plant pathogen) to evaluate growth inhibition.

Wild type, ΔII2523, Δcluster 15, and ΔII2523 Δcluster 15 equally inhibited growth of *B. subtilis* suggesting that malleipeptin is dispensable (Fig 11). Loss of cluster 2 or cluster 14 from either the wild-type or ΔII2523 backgrounds resulted in complete loss of *B. subtilis* growth inhibition indicating that the cryptic secondary metabolite produced by cluster 2 and syrbactin (cluster 14) are important in inhibiting *B. subtilis* growth (Fig 11). Interestingly, loss of cluster 11 in the wild-type Bp82 background could not inhibit *B. subtilis* growth, while the ΔII2523 Δcluster 11 mutant was still able to produce metabolites that interfered with *B. subtilis* growth (Fig 11).

**Figure 11.**
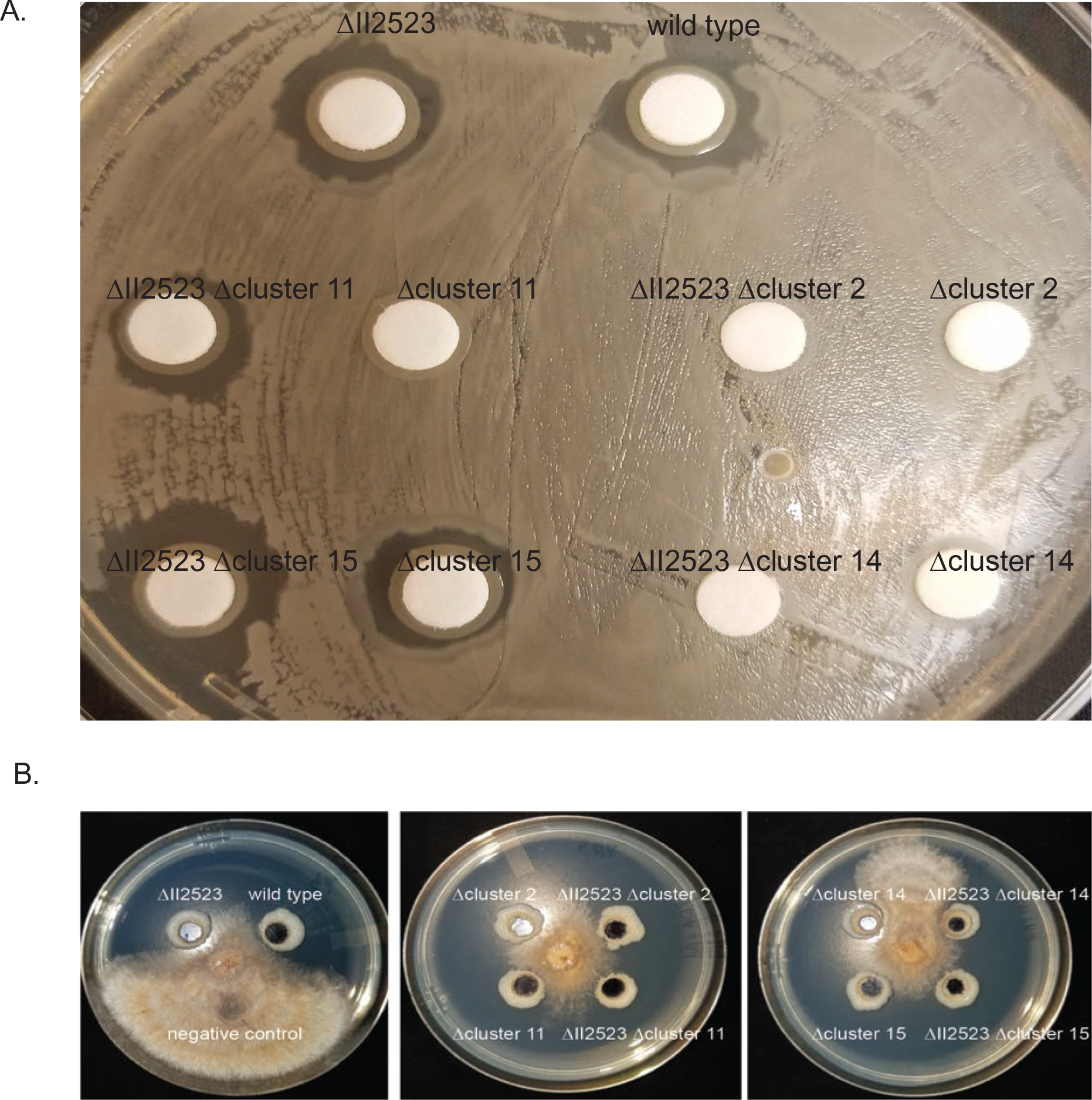
Growth inhibition of *B. subtilis* and *R. solani*. Zone of inhibition of *B. subtilis* and *R. solani* by the Bp82 wild type or Bp82 ΔII2523 NRPS deletion mutants. *B. subtilis* images were taken after 24h while *R. solani* images were taken after five days. All experiments were done at 28°C.

The contribution of these *B. pseudomallei* secondary metabolites was also evaluated in the context of a eukaryotic organism, *R. solani*, a soil-borne fungal pathogen of plants. All NRPS deletion mutants except for the cluster 14 (syrbactin) deletion mutant were able to strongly inhibit *R. solani* growth (Fig 11). These data suggest that syrbactin plays an integral role in limiting the growth of a eukaryotic organism like *R. solani* through an unknown mechanism. Purified syrbactin has been demonstrated to be a eukaryotic proteasome inhibitor in various eukaryotic cell lines (33); however, the mechanism of syrbactin inhibition of fungal growth is unknown.

These data indicate that cluster 14 (syrbactin) is important for inhibition of the growth of *B. subtilis* and *R. solani* while cluster 2 (uncharacterized NRPS) is important for specifically inhibiting *B. subtilis* in this study.

## Discussion

*B. pseudomallei* is a versatile organism that can transition from a saprophytic lifestyle residing in the soil to a pathogen of human and animal hosts. This adaptable organism can also form robust biofilms and produce an array of secondary metabolites in response to various environmental parameters which include exogenous nitrate (30) and quorum sensing signals (28). Correspondingly, the genome of *B. pseudomallei* encodes cryptic biosynthetic gene clusters (BGCs) that encode uncharacterized small molecules; although some metabolites have been characterized and produced using inducible promoters (14, 15). *Burkholderia* spp. have been reported to produce a diversity of metabolites that contribute to survival, adaptation, and interactions with other organisms (34).The vast majority of the environmental cues and signals that initiate the transition between lifestyles and regulate the production of metabolites are largely unknown in *B. pseudomallei*.

C-di-GMP has been shown to participate in controlling this switch in behavior or lifestyles at the level of a secondary messaging in many bacterial pathogens including *B. pseudomallei* (6, 35–37). To better understand the role of c-di-GMP in biofilm dynamics and the regulation of secondary metabolites in *B. pseudomallei*, we followed up on our previous studies of bacterial responses to nitrate and temperature (11, 30, 38). In this study, we generated in-frame deletion mutants of a select group of c-di-GMP genes and conducted an epistatic genetic analysis in the attenuated strain of *B. pseudomallei* Bp82 to unmask phenotypes that might have been previously hidden. The aim of this study was to further characterize the role of specific c-di-GMP genes in motility, biofilm formation, and colony morphology by generating double, triple, and quadruple mutants in c-di-GMP metabolic genes, which has not been previously done in *B. pseudomallei*. Since the temperature-dependent biofilm phenotypes of the II2523 transposon mutant were recapitulated with in-frame deletion of II2523, we performed RNA-seq analysis to identify genes that are important for the transition of planktonic cells to a biofilm mode of growth in *B. pseudomallei*. The RNA-seq study revealed a suite of polysaccharide, BGCs, and secretion system biosynthetic clusters that mediate wild-type and ΔII2523 biofilm formation at 28°C and 37°C.

The deletion of II2523 negatively affected biofilm formation at 28°C and 30°C resulting in decreased biofilm production as expected; however, this mutant hyperproduced biofilm at 37°C as previously observed in our transposon mutational analysis (11). These biofilm and swim phenotypes also correlated with decreased levels of c-di-GMP at 30°C and correspondingly higher levels of c-di-GMP at 37°C. Future studies will continue to address this unknown mechanism now that we have demonstrated these phenotypes using in-frame deletion mutants that can be used in the BSL2 laboratory.

Much of what we have previously learned concerning function and regulation of these cryptic metabolites is gleaned from genomic comparisons of the foundational members of the *Burkholderia pseudomallei* complex (Bpc), which are comprised of *B. pseudomallei*, *B. mallei*, and *B. thailandensis*. Genomic analyses of Bpc organisms provide clues to the functional role of cryptic molecules given that these closely related species have divergent lifestyles and provide the opportunity to identify the key factors that contribute to survival in the different environments and hosts that they can inhabit.

*B. pseudomallei*, which is the causative agent of melioidosis in humans, is an environmental saprophyte found in soils and surface waters in endemic regions (39), and can also cause disease in a variety of mammalian hosts, birds, and reptiles (40) and can survive in amoeba (41). In contrast *B. mallei* causes glanders in equids and is not found outside of mammalian hosts which is believed to be due to evolution through genome reduction and rearrangement from a single strain of *B. pseudomallei* (42)*. B. thailandensis* is a nonpathogenic relative of *B. pseudomallei* that is also a soil dwelling saprophyte (43, 44). For this study we focused on the *B. pseudomallei* 1026b genome, which is predicted to encode 15 NRPS/PKS BGCs, seven of which are not conserved in *B. mallei* (the genome-reduced relative of *B. pseudomallei*) and four of these BGCs are not conserved in *B. thailandensis* (14). The differences in the conservation of BGCs between these closely related species affords the opportunity to evaluate the contribution of these metabolites to survival in different niches (45).

In our RNA-seq data set under biofilm inducing conditions, BGCs 2, 3, 12, and 13 were differentially regulated, which were also and previously shown to be regulated by quorum sensing in planktonic growth conditions (28). However, very little is known about the functional role of these secondary metabolites in *B. pseudomallei*. In *B. thailandensis,* a malleilactone (BGC 8) mutant was significantly less virulent than wild-type *B. thailandensis* in a *C. elegans* killing model and did not inhibit *D. discoideum* from forming fruiting bodies (46). In *B. pseudomallei* K96243, a glidobactin (syrbactin) mutant was more susceptible to killing by human neutrophils. Interestingly, the glidobactin mutant was more lethal than the wild type at a high titer, while at low titer the survival between the mutant and wild type was comparable (16). This is in contrast to Biggins and colleagues that demonstrated that a syrbactin mutant in the *B. pseudomallei* 1026b background was highly attenuated in a mouse model (14). These contradictory results observed between the syrbactin mutants may be the result of the different genetic background of the strains used in these studies and/or the experimental parameters that were utilized.

The function of the various *B. pseudomallei* secondary metabolites encoded by the BGCs evaluated in these studies and their contribution to biofilm formation has yet to be fully elucidated. In this study, we have shown BGCs 2 and 11 have significant contributions to biofilm formation at 37°C. BGCs 2, 11, 14, and 15 also have significant contributions to biofilm formation at 28°C where biofilm formation would provide a competitive advantage for survival in the environment. The Bpc group has recently undergone a proposed expansion to include *B. oklahomensis*, *B. humptydooensis, Burkholderia mayonis* sp. nov. and *Burkholderia savannae* sp. nov. in addition to *B. pseudomallei*, *B. mallei*, and *B. thailandensis* based on whole genome sequence analyses (42, 47). This expansion will increase our ability to further resolve the function and association of specific BGCs with organisms that inhabit different niches. These characterized BGCs could also provide better resolution for diagnostics based on genetic and metabolic markers.

The results of these studies illustrate the potential of how this system could be used to evaluate the secreted polysaccharides and secondary metabolites produced by *B. pseudomallei* as observed by the effects of mutations in key BGCs have on biofilm formation and colony morphology. Future studies will evaluate the role of these molecules in stabilizing and protecting the biofilm matrix from degradation. Other biofilm producing strains like *Pseudomonas aeruginosa* are known to have protein structural components that fortify the biofilm matrix (21) that are regulated by c-di-GMP, and a protease inhibitor protein that protects the biofilm matrix from proteolytic attack (48). Biggins et al. proposed that the malleipeptins (cluster 15) could function as biosurfactants/biofilm modulators and the syrbactin-type proteasome inhibitors (cluster 14) were representative of the small-molecules that have gone unnoticed in *B. pseudomallei*. Based on the results of the colony morphology and biofilm assays presented it is visible that the molecules encoded by these BGCs alter the surface of these bacterial communities and manipulate biofilm dynamics.

Future studies will focus on the characterization of BGC 12 and 16 as this cluster was upregulated in the ΔII2523 at 28°C and is likely important for bacterial competition and protection from predation during *B. pseudomallei* biofilm growth. The genetic backgrounds and environmental growth conditions identified in these studies will allow further chemical characterization of these cryptic metabolites and their role in growth, survival, competition, and infection of hosts.

## Methods

### Bacteria growth, mutants, and complementation

*B. pseudomallei* Bp82 (BSL3 select agent excluded strain) was grown in LB supplemented with 80 µg/mL adenine (LB+Ad80) (13). *B. pseudomallei* 1026b (select agent) was grown in LB and handled in a BSL3 laboratory. Generation of in-frame deletion mutants of Bp1026b_I2284, Bp1026b_I2285, Bp1026b_I2284-Bp1026b_I2285, Bp1026b_II0885, Bp1026b_II2523 was accomplished by allelic exchange as previously described (49). SOEing PCR was used to amplify approximately 1kB of flanking sequence on both sides of the gene(s) or a synthesized fragment for *bce-II* (Genescript). Deletion constructs for cluster 2, 11, 14, and 15 in pEXKm5 were kindly provided by D. Shazer (14). The resulting fragments were cloned into pEXKm5 (49) and electroporated into RHO3. RHO3 pEXKm5 overnight cultruess were grown in diaminophilic acid (DAP, 400 µg/mL) and kanamycin (35 µg/mL) and conjugated with Bp82. Merodiploids were selected on LB+DAP+1000 µg/mL kanamycin. Kanamycin resistant clones were re-struck onto LB+1000 µg/mL kanamycin 100 µg/mL +100 µg/mL 5-bromo-4-chloro-3-indolyl-β-D-glucuronide (X-gluc) plates at 37°C. The following morning, several blue colonies were used to inoculate YT (8 g/L tryptone and 5g/L yeast extract) broth containing 15% sucrose and were grown with shaking for several hours at 37°C and then plated onto YT (8 g/L tryptone, 5 g/L yeast extract, 15% sucrose) plates and 100 µg/mL X-gluc. White colonies were re-struck onto YT plates (15% sucrose and 100 µg/mL X-gluc) at 37°C and single colonies were patched onto LB plates with and without 1000 µg/mL kanamycin. Kanamycin-sensitive clones with presumed deletions were verified using internal and external primers to a gene within the cluster or flanking the cluster of interest. For complementation and conditional IPTG expression studies, Bp1026b_I2284 (*cdpA*), Bp1026b_I2285, Bp1026b_I2284-Bp1026b_I2285, Bp1026b_II0885, and Bp1026b_II2523 were PCR amplified from Bp82 genomic DNA using Phusion DNA polymerase (NEB) or Kapa HiFi polymerase (KapaBiosystems) using the primers (forward primers included a <u>ribosome binding sequence</u>) listed in (Supplemental Table 1). The corresponding fragments were ligated into the integration vector pUC18T-mini-Tn*7*T-km-LAC for expression in *B. pseudomallei* or *P. aeruginosa* (50). Clones were sequenced to confirm that no mutations had been introduced. The resulting vectors were electroporated into DH5α cells. Plasmids were isolated and electroporated in *E. coli* RHO3. RHO3 cells harboring the pUC18T-mini-Tn*7*T-KM-LAC constructs were grown in LB with DAP 400 µg/mL and kanamycin 35 µg/mL and mixed with RHO3 pTNS3 in Bp82 triparental matings. Complemented mutants and empty vector controls were selected on LB plates with 1000 µg/mL kanamycin. A final of 1mM IPTG was used for gene expression. Site-directed mutagenesis was done according to manufacturer’s recommendation using a QuikChange Lightning kit (Agilent Genomics) as we have previously described (11) using the oligonucleotides listed in Supplemental Table 1. All plasmids were confirmed via sequencing.

### Static biofilm and swim motility assays

Biofilm and swim motility assays were done as previously described by (11) with the addition of adenine in the media. *P. aeruginosa* strains (PAO1 or PAO1 Δ*wspF*) conditionally expressing Bp1026b_I2284 (*cdpA*), Bp1026b_I2285, Bp1026b_I2284-I2285, Bp1026b_II0885, Bp1026b_II2523, or empty vector (pUC18T-mini-Tn*7*T-KM-LAC) were grown overnight in LB (kanamycin 300 µg/mL where appropriate).

### C-di-GMP measurements

C-di-GMP extractions and quantification were done as described by Mangalea et al. (38) with modifications using the BSL-3 parental strain 1026b and an in-frame deletion mutant ΔII2523. Briefly, overnight cultures were grown in LB at 37°C with shaking at 250 rpm, diluted 1:50 in M9 Salts minimal medium, and grown statically for 18 h at either 30°C or 37°C. Extractions were performed using a chilled matrix buffer solution of acetonitrile/LC-MS methanol/LC-MS H_2_O/1% formic acid, supplemented with 100 nM of 2-chloro-adenosine-5’-*O*-monophsphate (2-Cl-5’-AMP, Axxora, LLC) for internal standardization. Final absolute nucleotide concentrations were normalized to total protein concentrations using the Pierce 660 nm Protein Assay (Thermo Scientific). Extraction experiments were repeated on separate days using two biological replicate cultures for each strain and temperature condition, with three technical replicates each. Statistical significance was calculated using an unpaired Student’s *t*-test in GraphPad Prism (v7) with the Bonferroni-Sidak correction for multiple comparisons.

### RNA isolation and RNA sequencing

Total RNA was collected from pellicle biofilms formed at the air-liquid interface from three technical replicates per biological sample grown in six well Costar plates for 24h at either 28°C or 37°C. Samples in which pellicles were inhibited due to treatment were collected from bacterial cells at the bottom and sides of the Costar plate. Each plate was seeded with three mutant replicates and three wild-type replicates for each condition tested. 1.5 mL of samples were collected and spun at 12,000 rpm for 2 minutes and resuspended in 350 µL RNA Protect Bacterial Reagent (Qiagen). Samples were kept on ice from this point forward. Samples were spun at 5000 x g for 10 minutes and supernatant was discarded before resuspension in 1.5 mL QIAzol reagent (Qiagen). Screw-cap tubes were prepared for each sample by adding ∼250 µL of sterile beads and the QIAzol suspension mixture was added and incubated for 5 minutes at room temperature. Sample tubes were transferred back to ice before 3 rounds of 60 second bead beating on the TissueLyser2 (Qiagen). Samples were incubated at room temperature for five minutes before 200 µL of chloroform (Fisher Scientific) was added and samples were vortexed for five seconds before another room temperature incubation for five minutes. Samples were spun at 10,000 rcf for 10 minutes and 500 µL of the aqueous phase was removed and transferred to separate tubes containing 500 µL of molecular grade 70% ethanol. RNA was subsequently extracted from the samples using the RNeasy Kit (Qiagen), where the three technical replicates were pooled onto a single RNeasy column. Genomic DNA was removed using DNaseI via two rounds of treatment using the TURBO DNA-free Kit (Ambion). Ribosomal RNA was removed using RiboZero rRNA removal reagents for Gram-negative bacteria (Illumina). Libraries were prepared following the RNA-seq sample work flow using the Scriptseq Complete kit for bacteria and indexed with unique Scriptseq Index PCR primers (Illumina).

Samples were analyzed on a Tapestation using HS D1000 tapes and reagents (Agilent) to determine average sizes and concentrations of the libraries. Size and molarity estimates were used to pool all libraries in equimolar concentrations. Final quality control and qPCR analyses were completed at the Colorado State University NGS Core Facility. A NextSeq run was completed on the pooled libraries using the NextSeq 500 hi-output v2 75-cycle kit and Buffer Cartridge (Illumina). Sequence files were downloaded from the NGS server, de-multiplexed according to index primers, and converted to FastQ files before initial quality control using FastQC (version 0.10.1) (51). Adapter sequences were trimmed using Trimmomatic (version 0.35) (52) before another quality control round using FastQC. Bowtie2 (53) was used to align sequencing reads to the reference genome GCF_000260515.1 and ASM26051v1 assembly (NCBI) and TopHat (54) was used for transcriptome mapping. HTseq (version 0.11.0) (55) was used to count accepted hits before the DEseq2 (version 1.20.0) (27) package was employed in R for comprehensive differential expression analysis. Raw read count coverage values were used to compare the differential gene expression between temperature treatments, mutants, and untreated controls. Using a negative binomial distribution to estimate variance and a Bayesian approach for variance shrinkage, the DEseq2 package produced logarithmic fold-change values between the conditions tested. Wald tests were used to calculate p-value and the Benjamini-Hochberg multiple testing correction was used to correct for the false discovery rate.

### Gene expression by quantitative real-time PCR

RNA was isolated using identical conditions to the samples used for RNA sequencing. Briefly, genomic material was isolated from static cultures grown for 24 h at the temperature indicated, using RNAprotect Bacterial Reagent (Qiagen) and QIAzol Lysis Reagent (Qiagen) before purification using RNeasy Mini Kit columns (Qiagen). Total RNA samples were treated with DNase I (Ambion/Life Technologies) twice, and cDNA was synthesized using Transcriptor First Strand cDNA Synthesis Kit (Roche) following the protocol recommended by the manufacturer. Primers were designed using the PrimerQuest tool (Integrated DNA Technologies). The 23S rRNA subunit was used for normalization in all experiments, using previously published primers (56). qRT-PCR experiments were performed on a LightCycler 480 instrument (Roche), in 96-well plates, using technical triplicates, with the reference gene present on all plates. Three biological replicates were included in all analyses. As a control for genomic DNA contamination in the cDNA samples, a set of control samples containing no reverse transcriptase enzyme were used in all experiments. Data were analyzed using the Pfaffl method (57), which considers the PCR efficiency percentage of all primer pairs including the reference gene.

### Colony morphology

Overnight cultures of *B. pseudomallei* strains were grown in LB. Then 3 µL were spotted on either LB, NAP-A, or YEM. Plates were incubated at either 28°C or 37°C for three days. Colony morphology images were taken with a Leica MZ95 microscope.

### Growth inhibition assay of Bacillus subtilis and Rhizoctonia solani

*B. pseudomallei* and *B. subtilis* 3610 overnight cultures were grown at 37°C with shaking (250 rpm). *B. subtilis* was diluted to a final OD_600_ of 0.1 and 100 µL of diluted culture was spread onto (LB+Ad80) plates until completely dry. Sterile filter discs (Remel, Lenexa, Kansas) were placed with sterile forceps onto plates and 20 µL *B. pseudomallei* cultures, diluted to OD_600_ 0.5, were dispensed onto the filter discs. Plates were allowed to incubate at either 28°C overnight. *R. solani* AG2-2 IIIB R-9 (J. Leach, Colorado State University) was propagated on potato dextrose agar (PDA) and grown at 28°C. Agar plugs were removed from PDA + Ad80 and filled with 50 µL of overnight cultures of wild type or ΔII2523 NRPS mutants. Then an agar plug of 3-5 day old culture of *R. solani* R-9 was placed in the middle of the plate. Plates were incubated at 28°C for five days.

### Supplemental methods Reverse transcriptase PCR

Total RNA was isolated from exponential phase cultures (OD_600_ 0.7 – 0.8) grown in 3 mL LB cultures at 37°C shaking at 250 rpm. Following double treatment with DNaseI (Ambion/Life Technologies), 500 ng RNA was reverse transcribed using random hexamers and the Transcriptor First Strand cDNA Synthesis Kit (Roche) following the protocol recommended by the manufacturer. To analyze co-transcription of Bp1026b_I2284 (*cdpA*) and Bp1026b_I2285, the cDNA was amplified using the following PCR primers: 5’-GACGCGATCAAGCTGACGGGC-3’ and 5’-CGAACGCCATGTACATCGCCG-3’. This primer set generated a 719 bp amplicon spanning the intergenic region of *cdpA* and I2285. A high-fidelity KAPA polymerase was used with 5x GC buffer with the following cycle conditions: pre-incubation at 95°C for 30 s, followed by 25 cycles of denaturation at 98°C for 20 s, annealing at 62°C for 15 s, extension at 72°C for 15 s, and a final incubation at 72°C for 1 min. 50 µL reactions were split into duplicates and visualized on an agarose gel. Bp82 genomic DNA was included as a positive control and cDNA from Δ*cdpA*-I2285 mutant was included as a negative control.

### Heterologous expression of *B. pseudomallei* c-di-GMP genes

Heterologous expression of *B. pseudomallei* c-di-GMP genes was done as described in the methods of this paper with the exception of triparental matings that were conducted with PAO1 and PAO1 Δ*wspF* as recipients.

## Acknowledgements

This work was supported by grant AI065357 from the National Institute of Allergy and Infectious Diseases, National Institutes of Health to the Rocky Mountain Regional Center of Excellence for Biodefense and Emerging Infectious Diseases Research, Colorado State University. We thank David DeShazer for providing pEXKm5 deletion constructs for clusters 2, 11, 14, and 15.

## Supplemental

**Fig S1.**
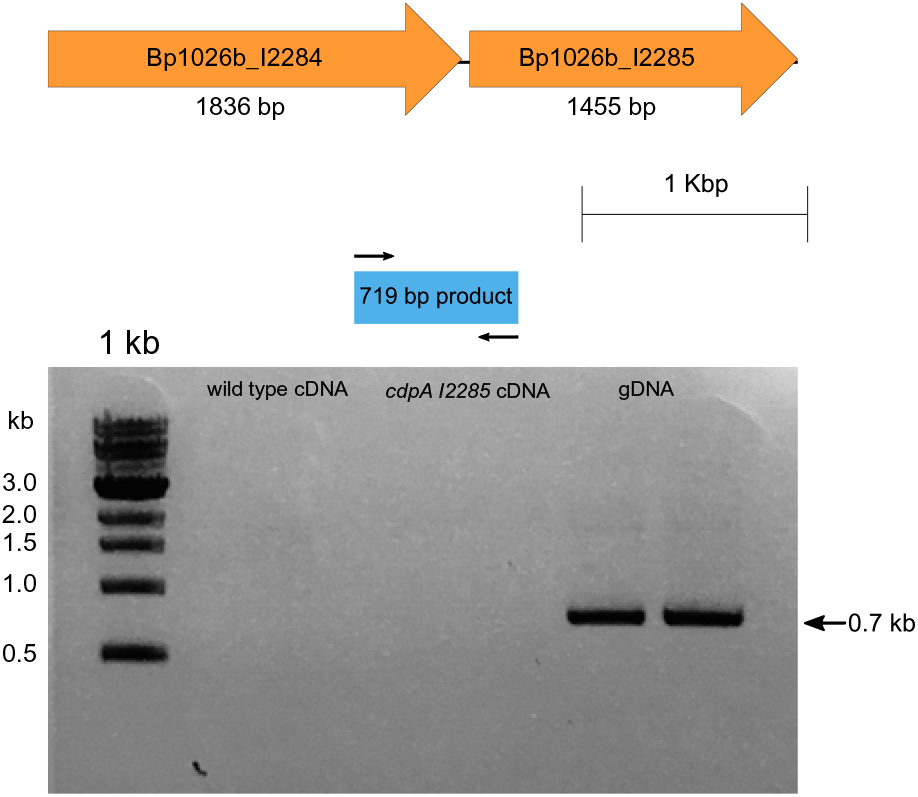
CdpA (I2284) and I2285 are not co-transcribed. cDNA from wild type, the Δ*cdpA-*I2285 double mutant, and wild-type genomic DNA were amplified to produce an expected product of 719bp. This product is only produced in genomic DNA where bands are visible for technical duplicate samples.

**Fig S2.**
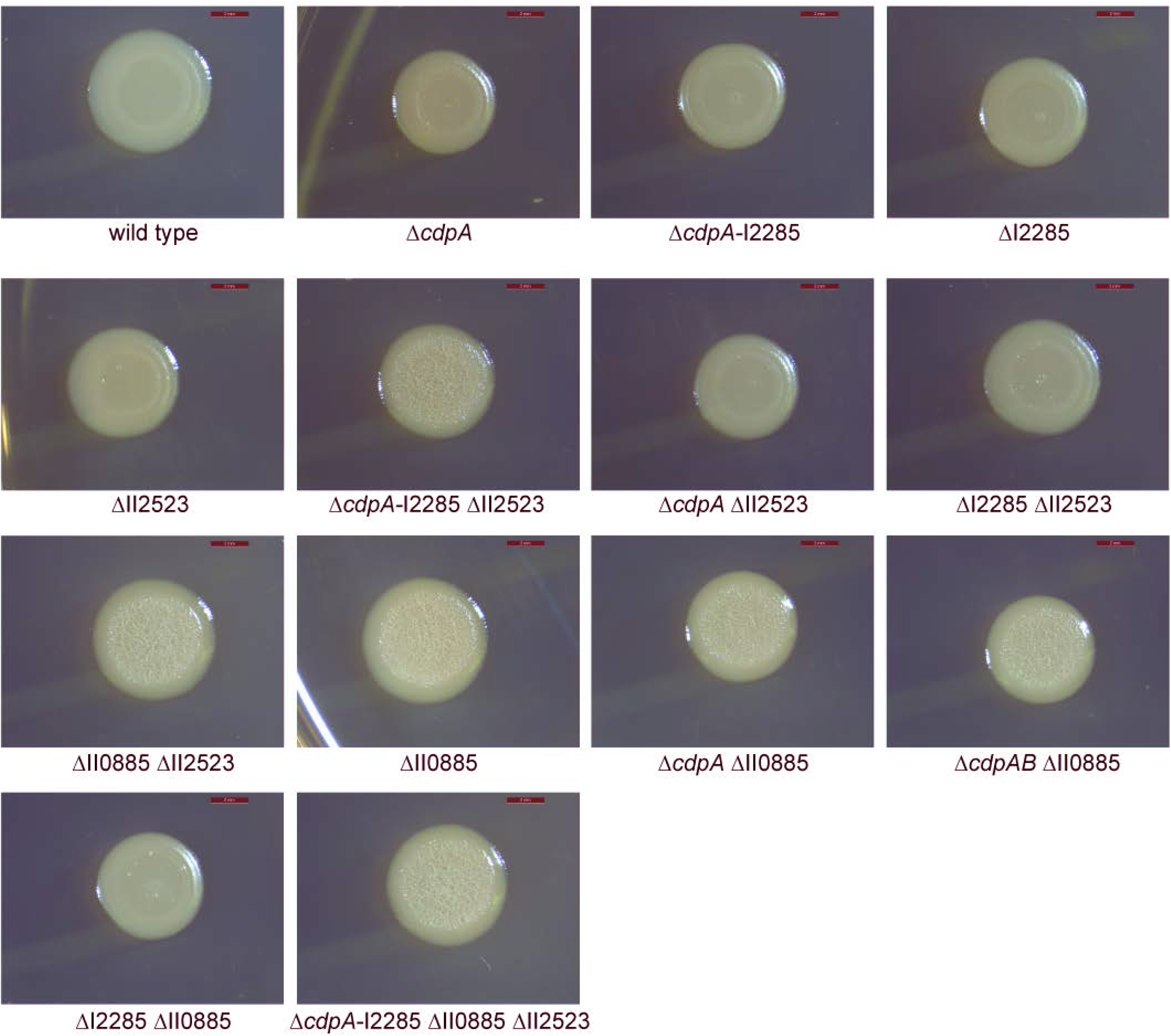
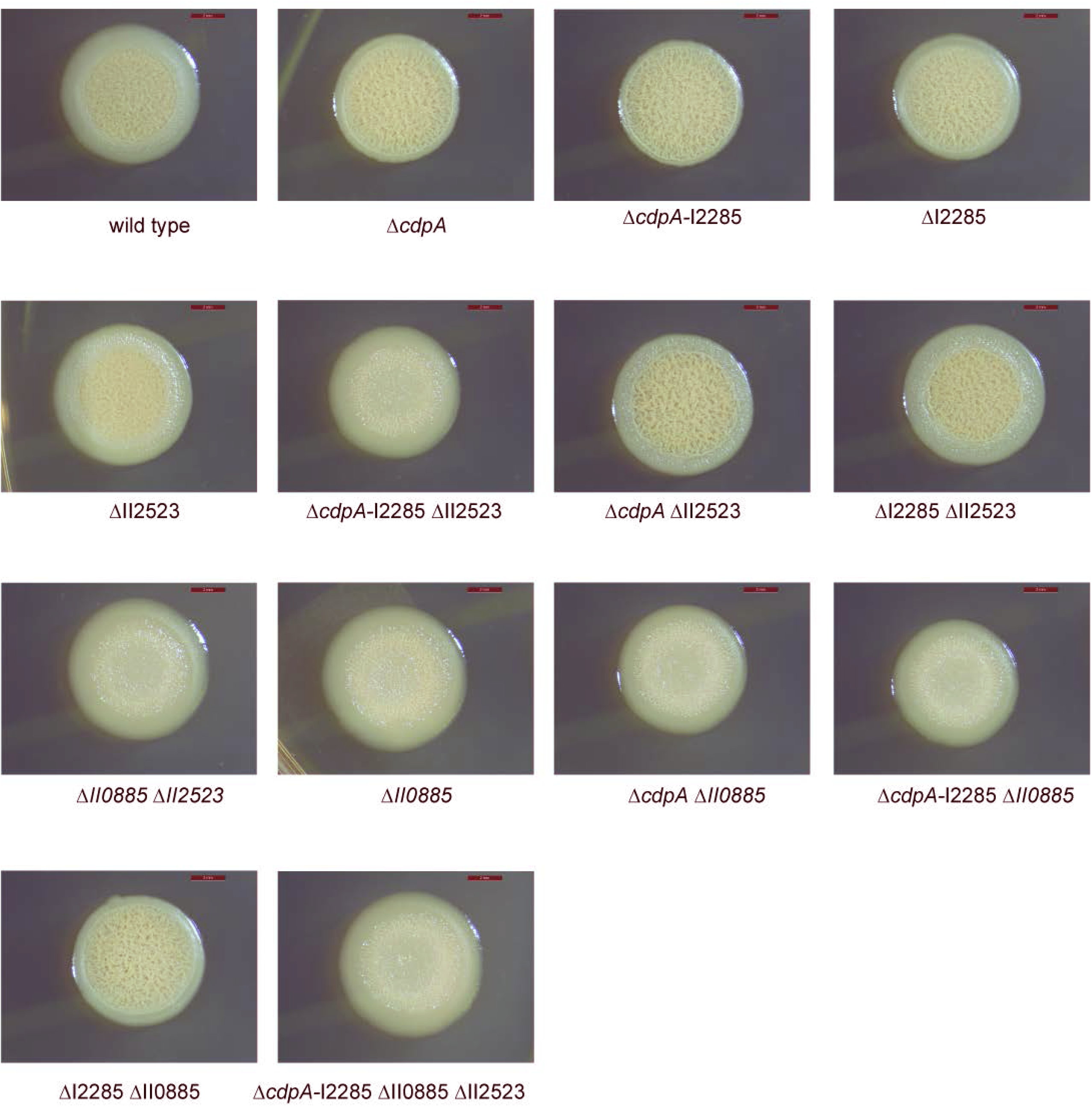
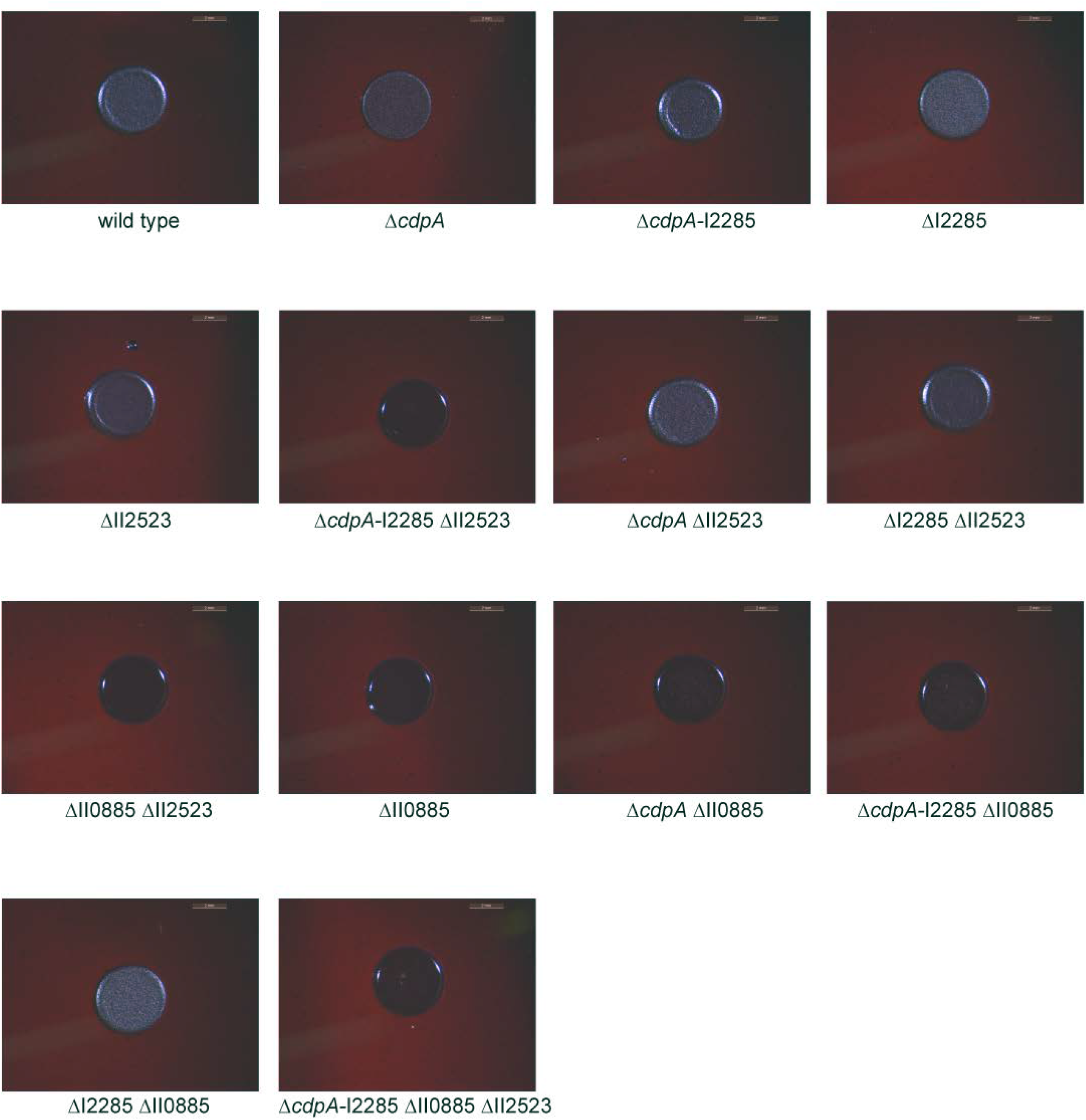
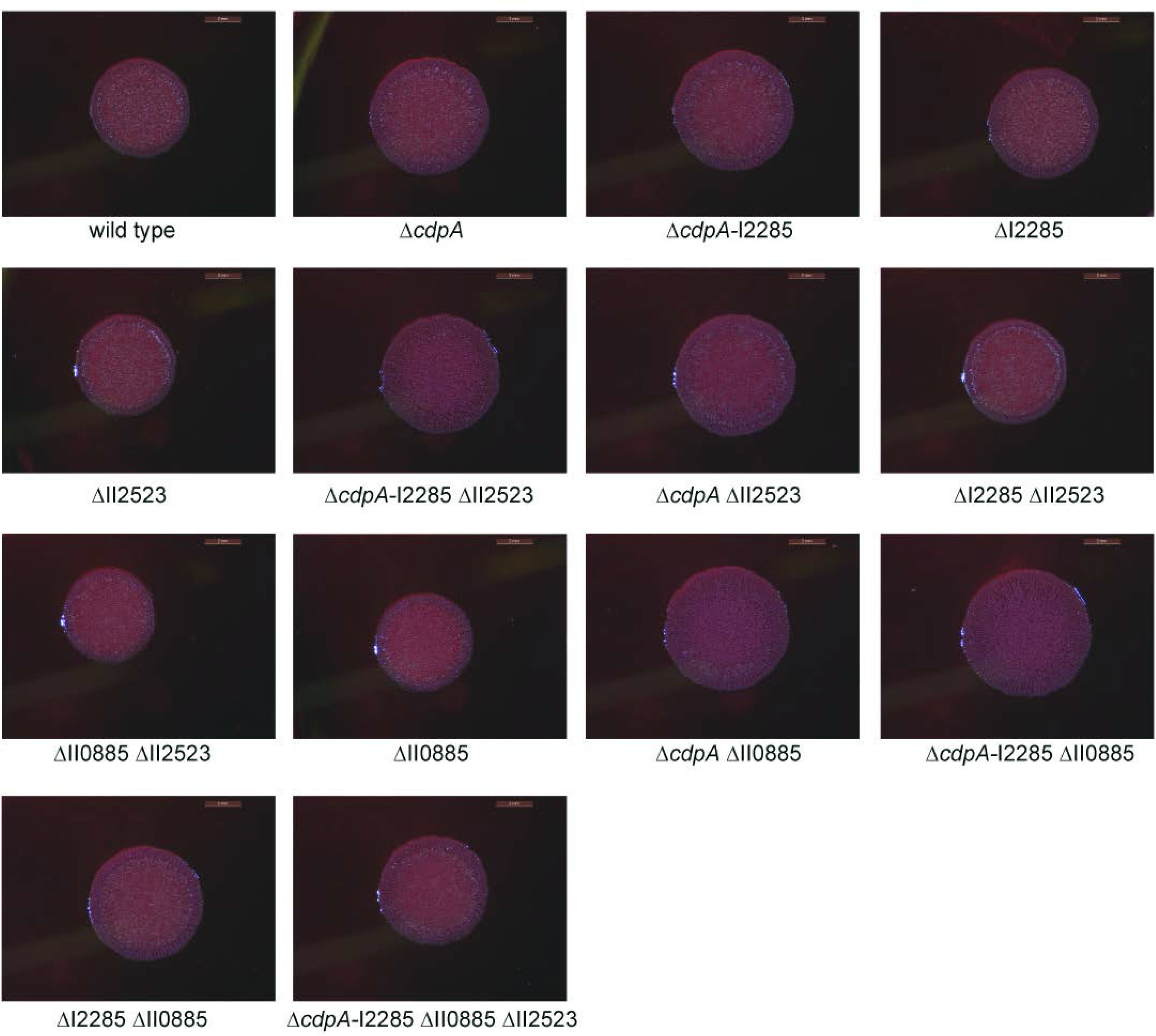
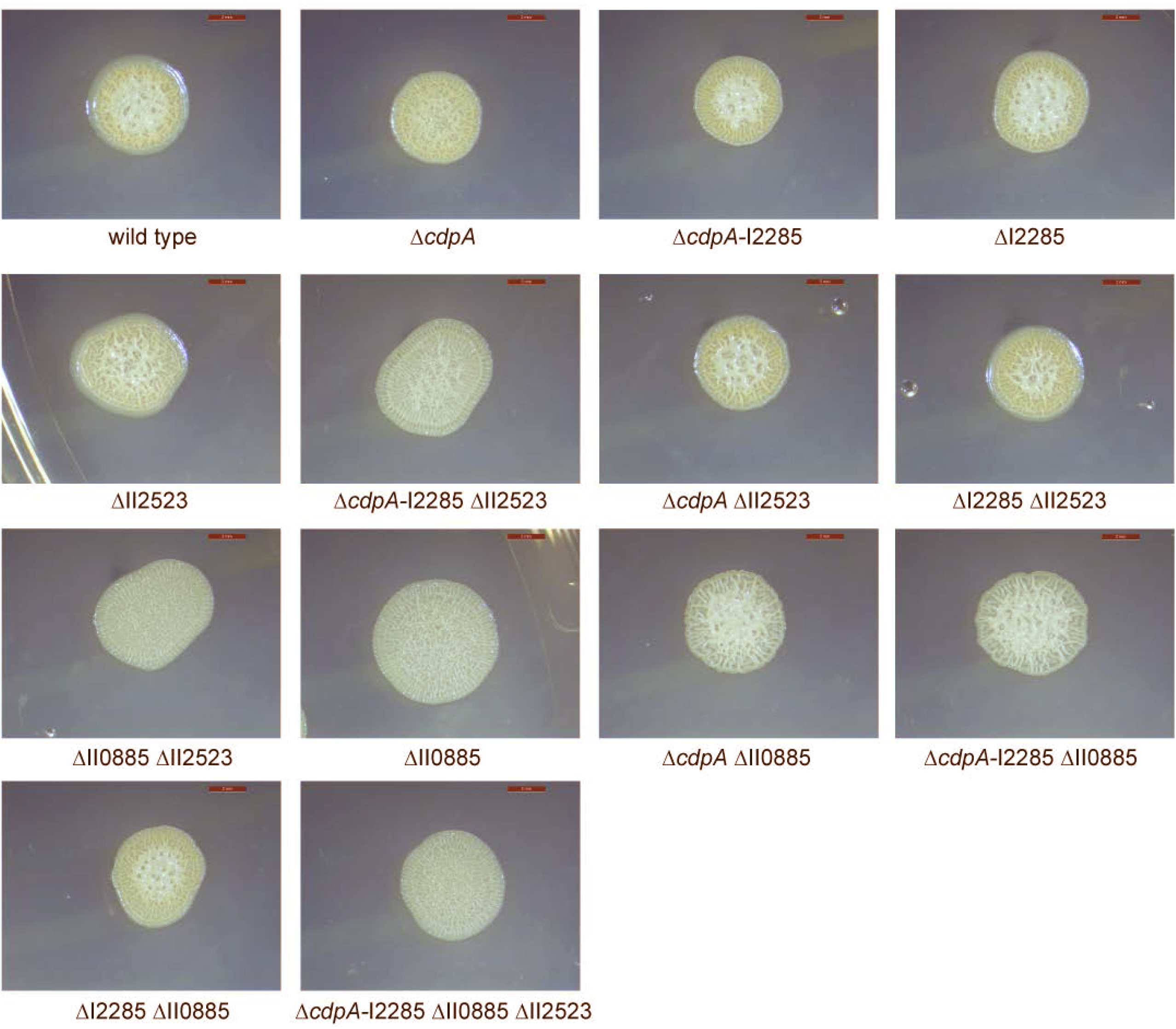
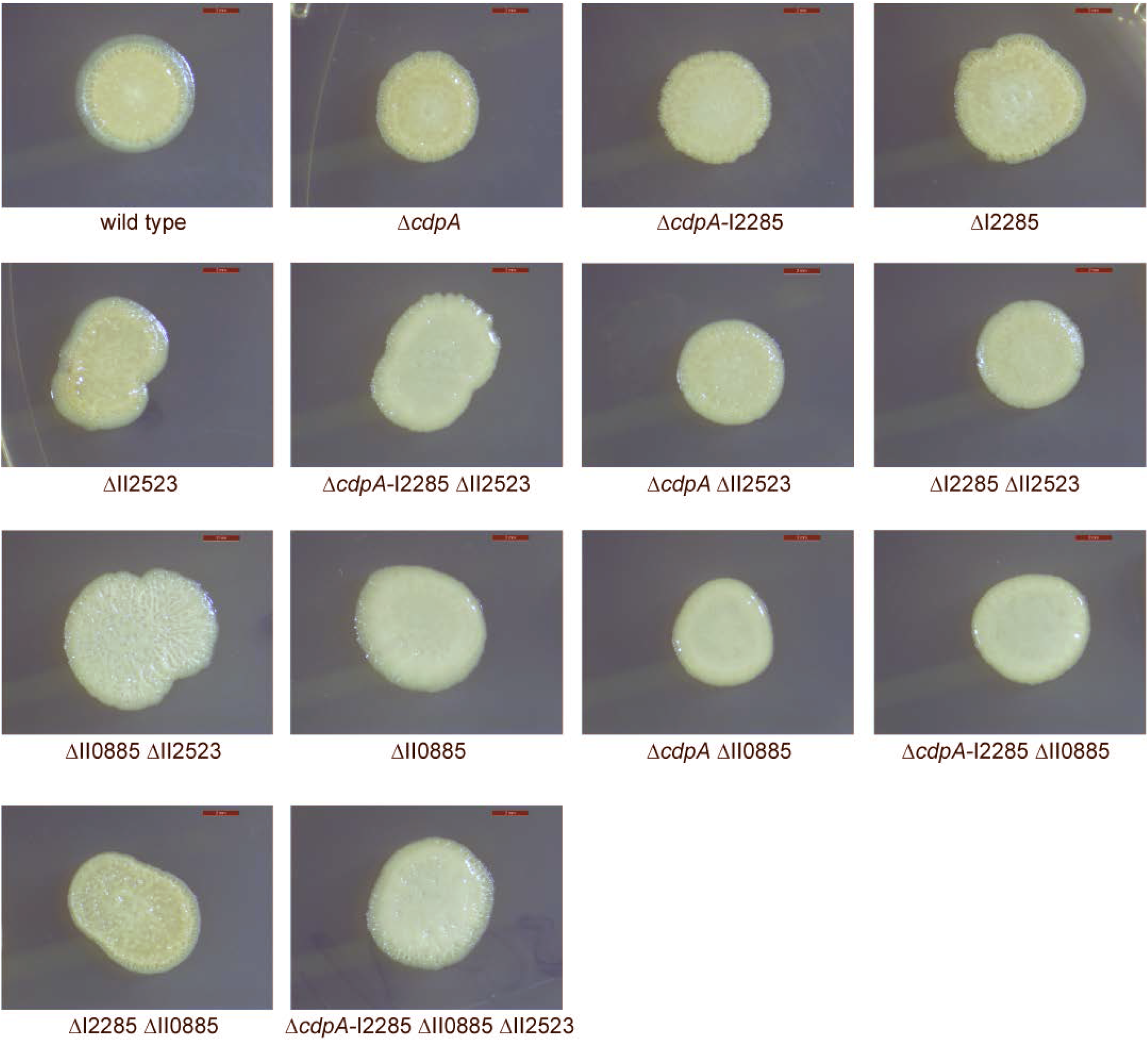
Colony morphology of c-di-GMP deletion mutants on LB at 28°C (A), LB at 37°C (B), NAP-A at 28°C (C), NAP-A at 37°C, YEM at 28°C (D), and YEM at 37°C (E). Images were taken at day three.

**Fig S3.**
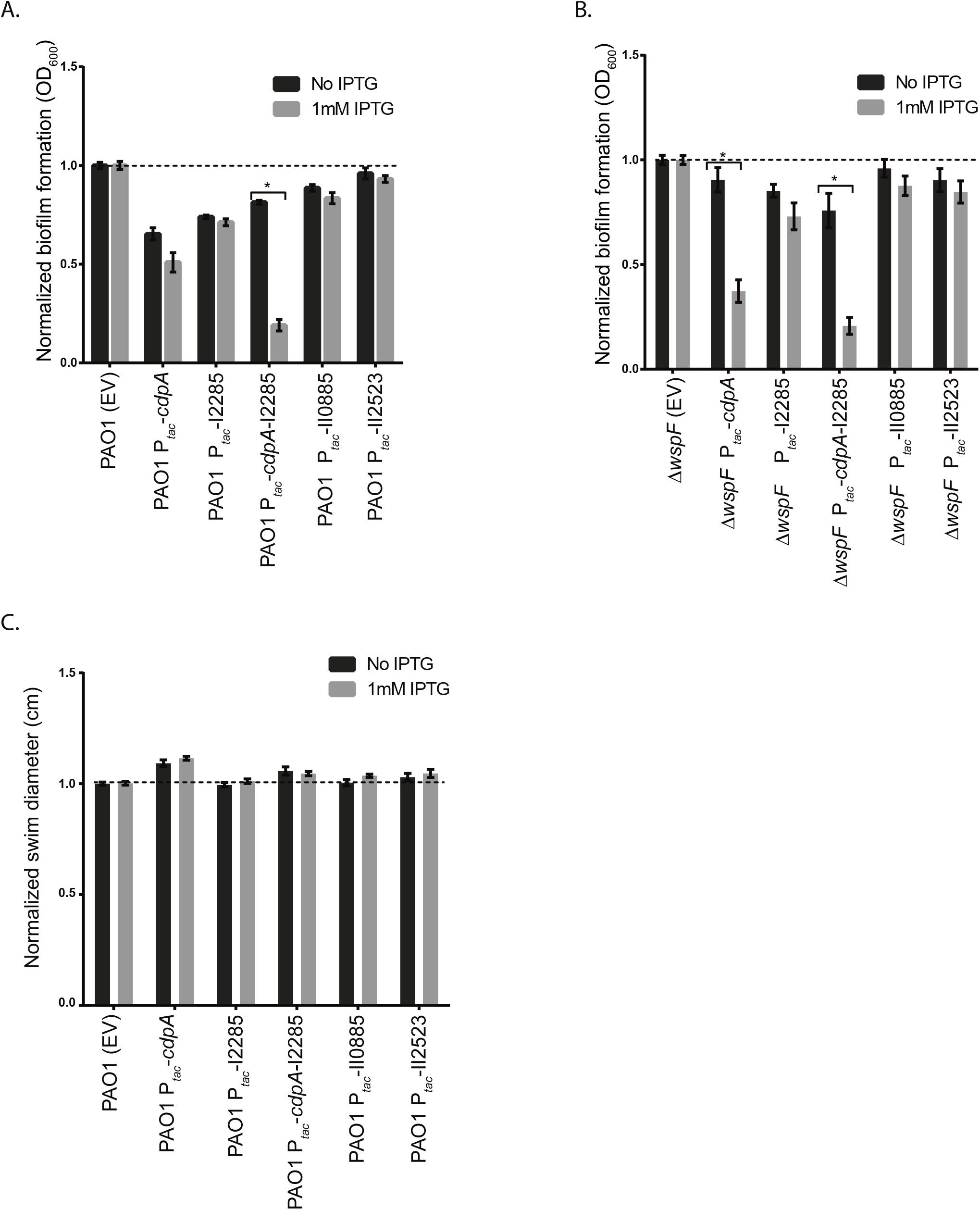
Biofilm formation and swimming motility of PAO1 and PAO1 Δ*wspF* strains heterologously expressing *cdpA,* I2285*, cdpA* I2285, II2523 and II0885. Biofilm formation PAO1 (A) and PAO1 Δ*wspF* (B). Swimming motility of PAO1 strains (C).

**Fig S4.**
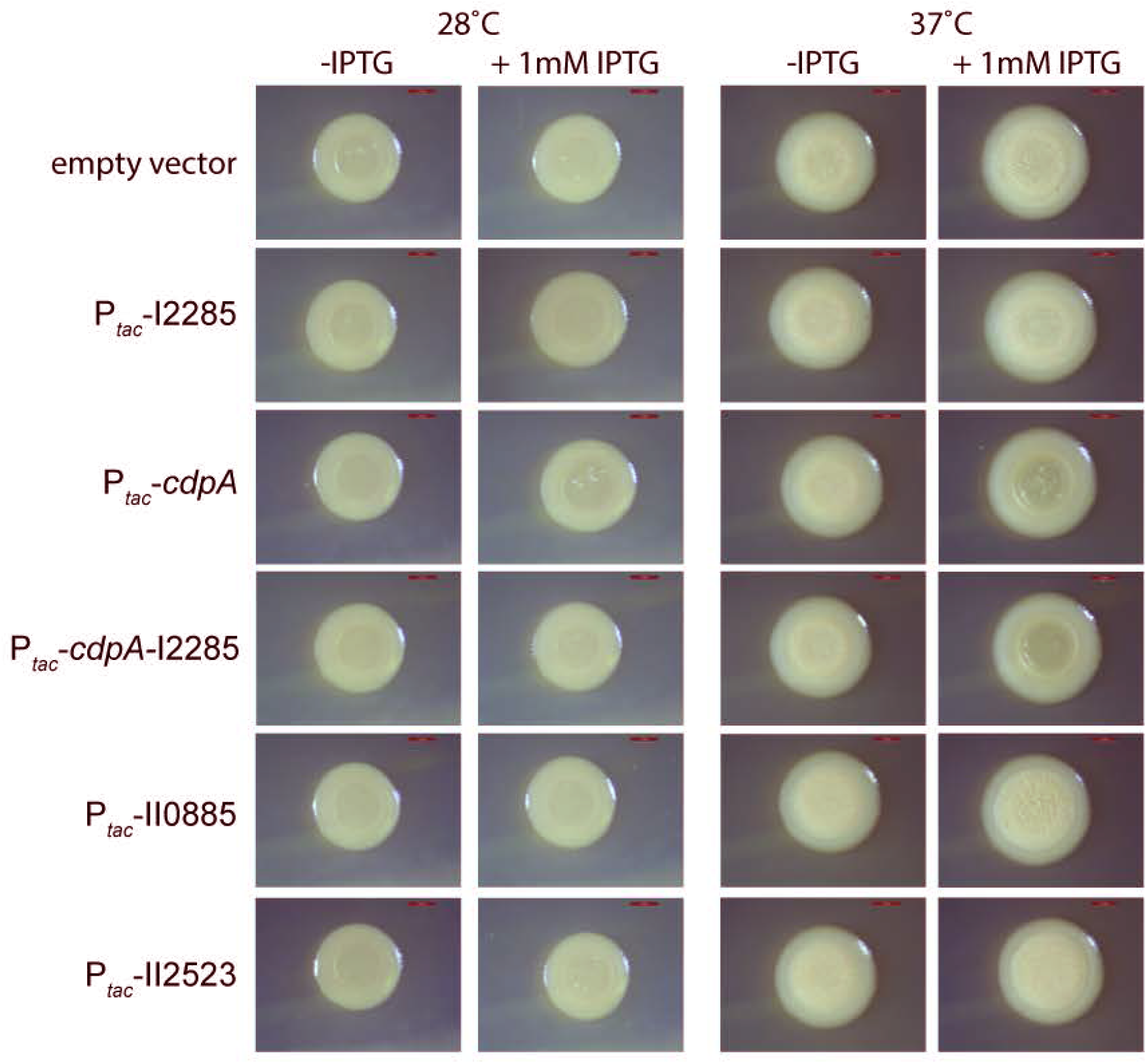
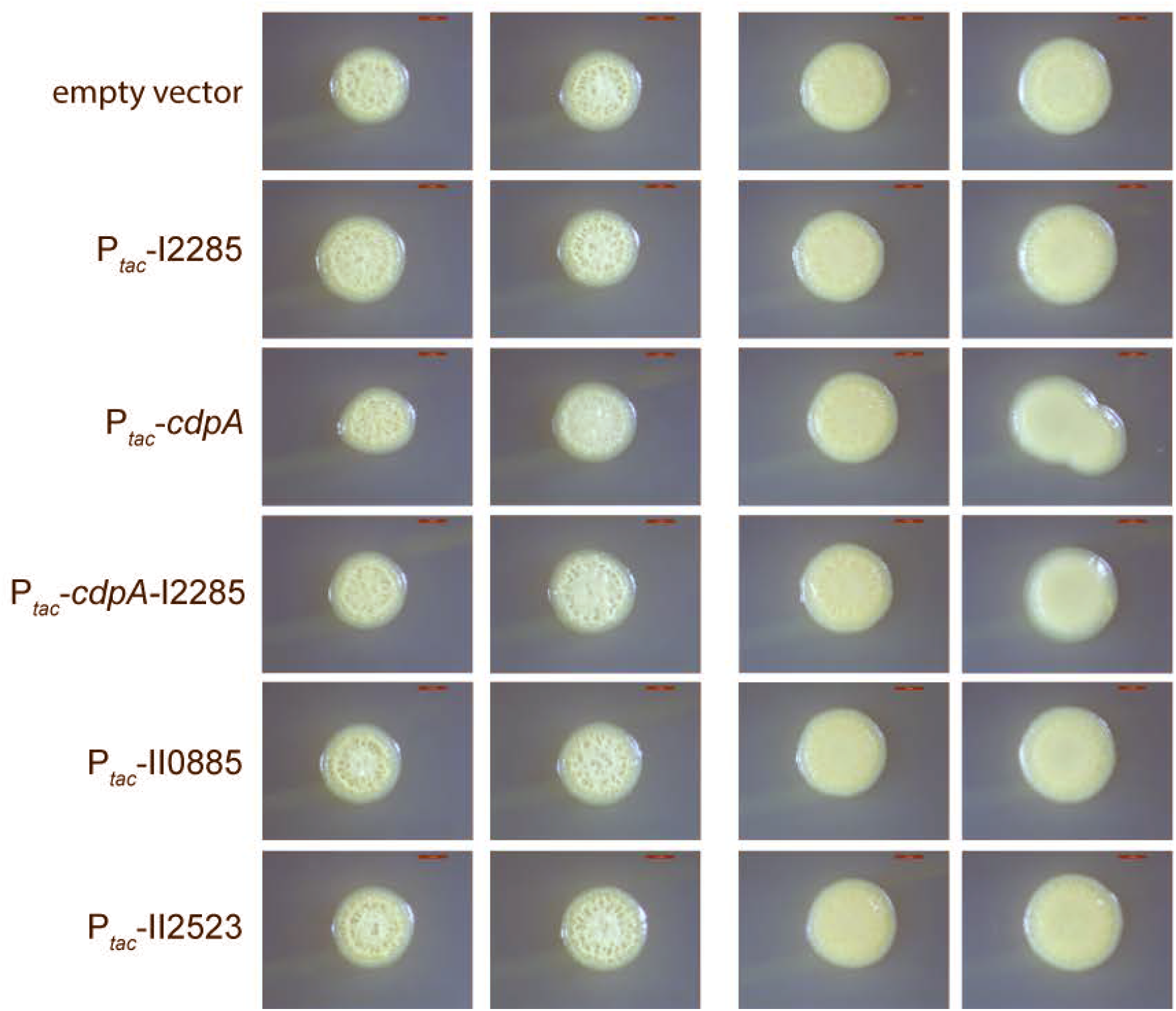
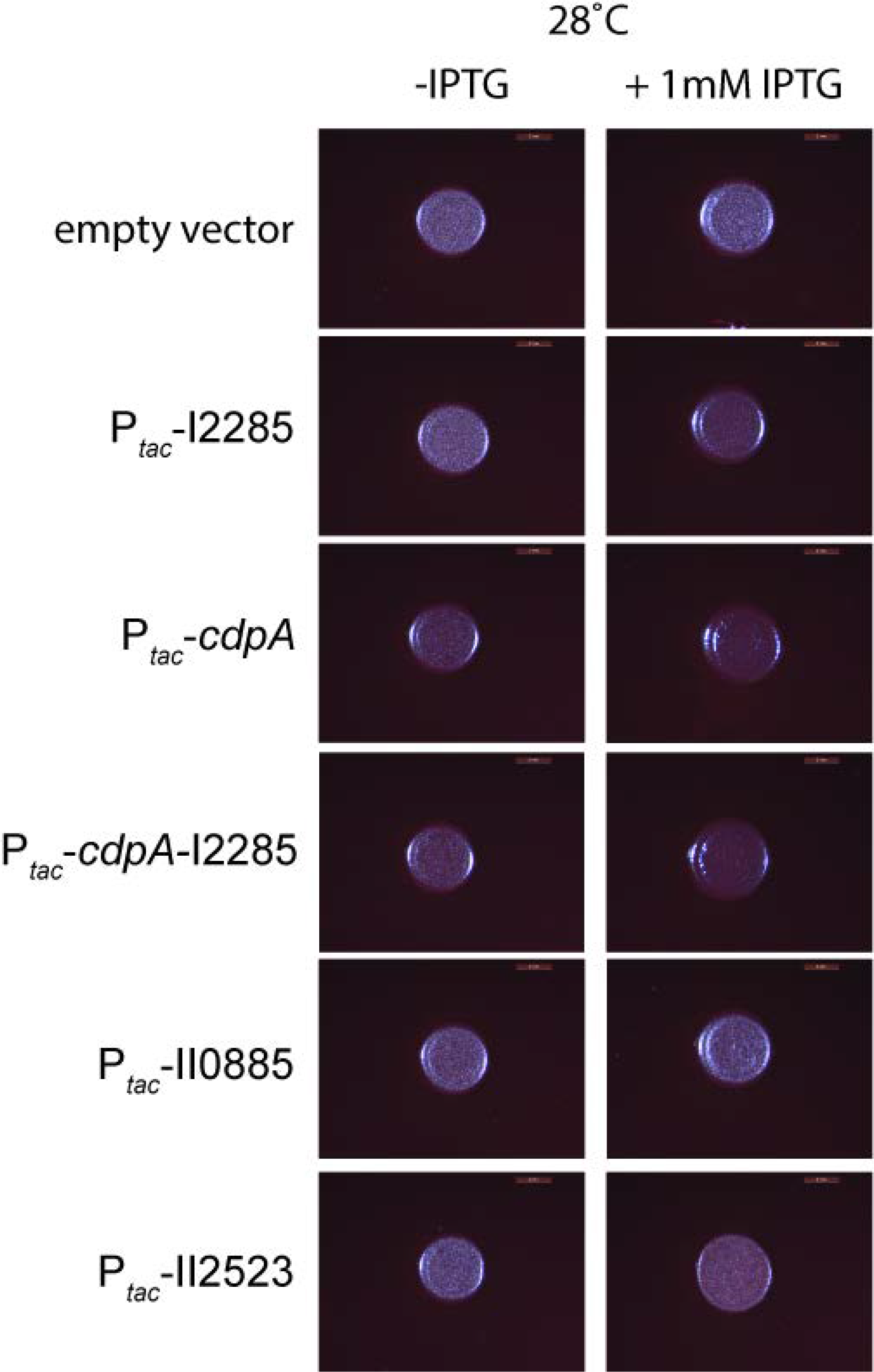
Inducible expression of c-di-GMP genes in wild-type *B. pseudomallei* Bp82. LB at 28°C and 37°C (A), YEM at 28°C and 37°C (B), and NAP-A at 28°C (C).

**Fig S5.**
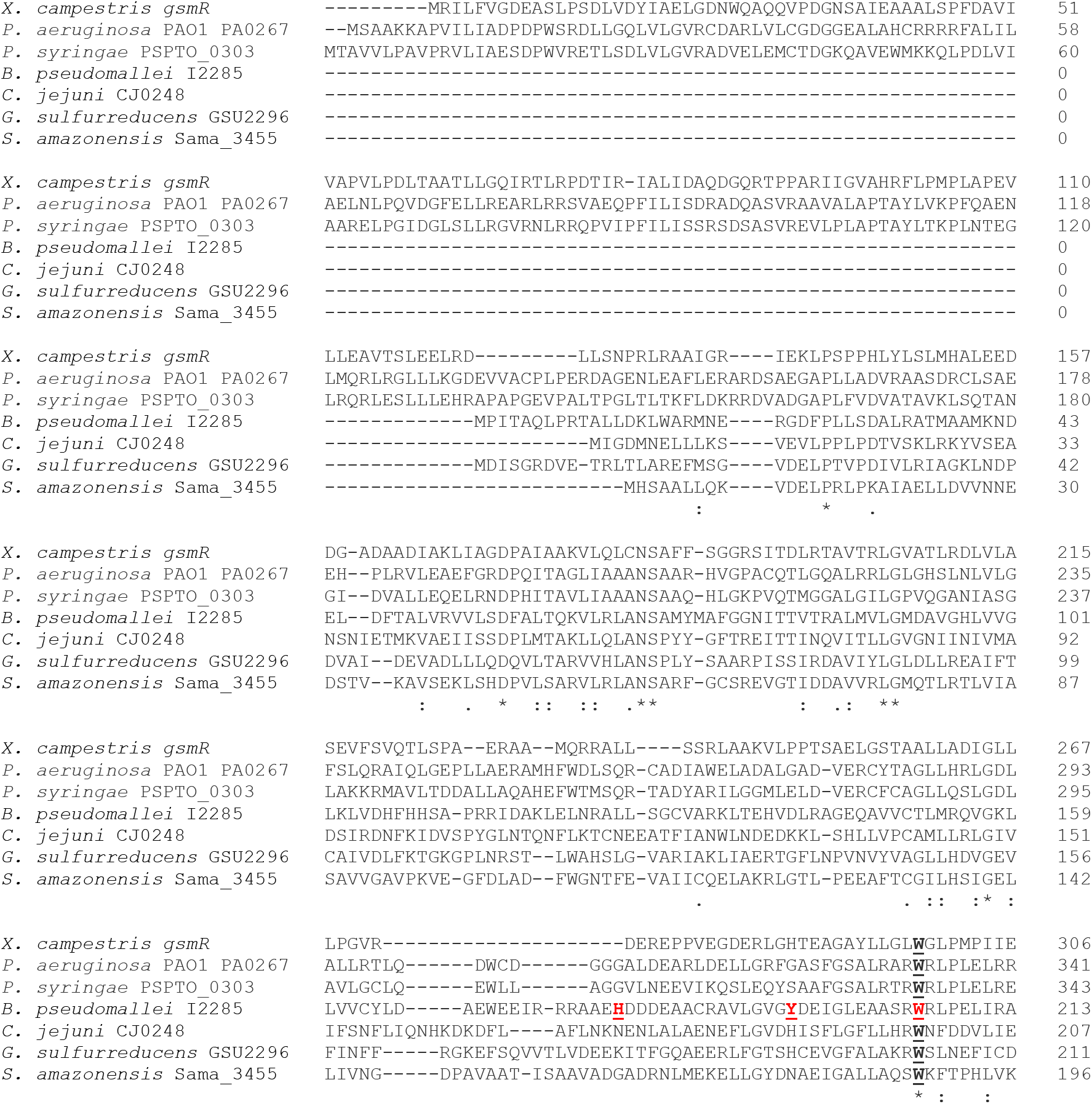
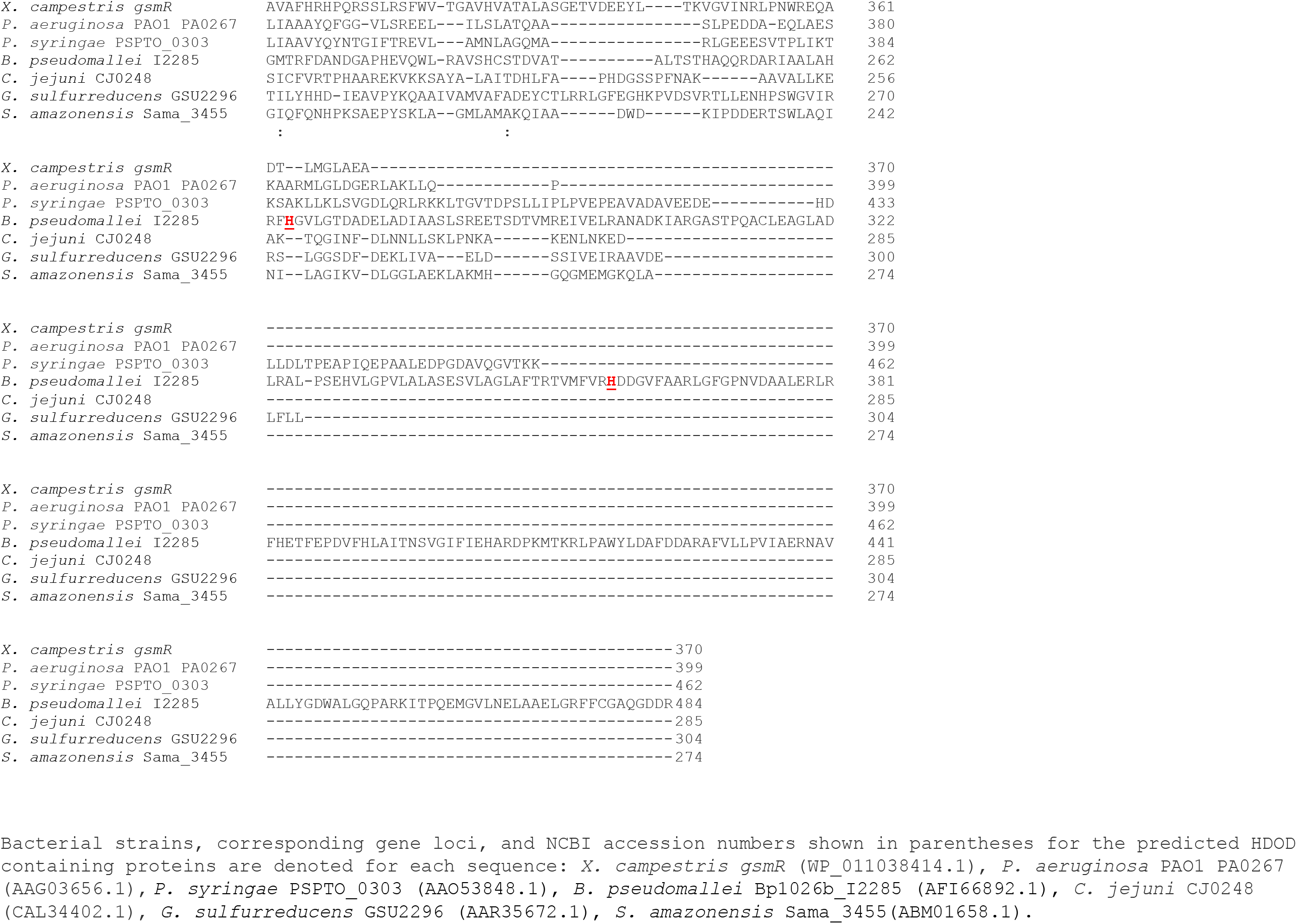
*B. pseudomallei* I2285 alignment with other predicted HDOD containing proteins from diverse bacterial strain. Protein alignment was done using clustal omega (Clustal 0 (1.2.4) multiple sequence alignment. *P. syringae* = *P. syringae* pv. tomato DC3000. Residues in red and underlined were targeted for site-directed mutagenesis.

**Fig S6.**
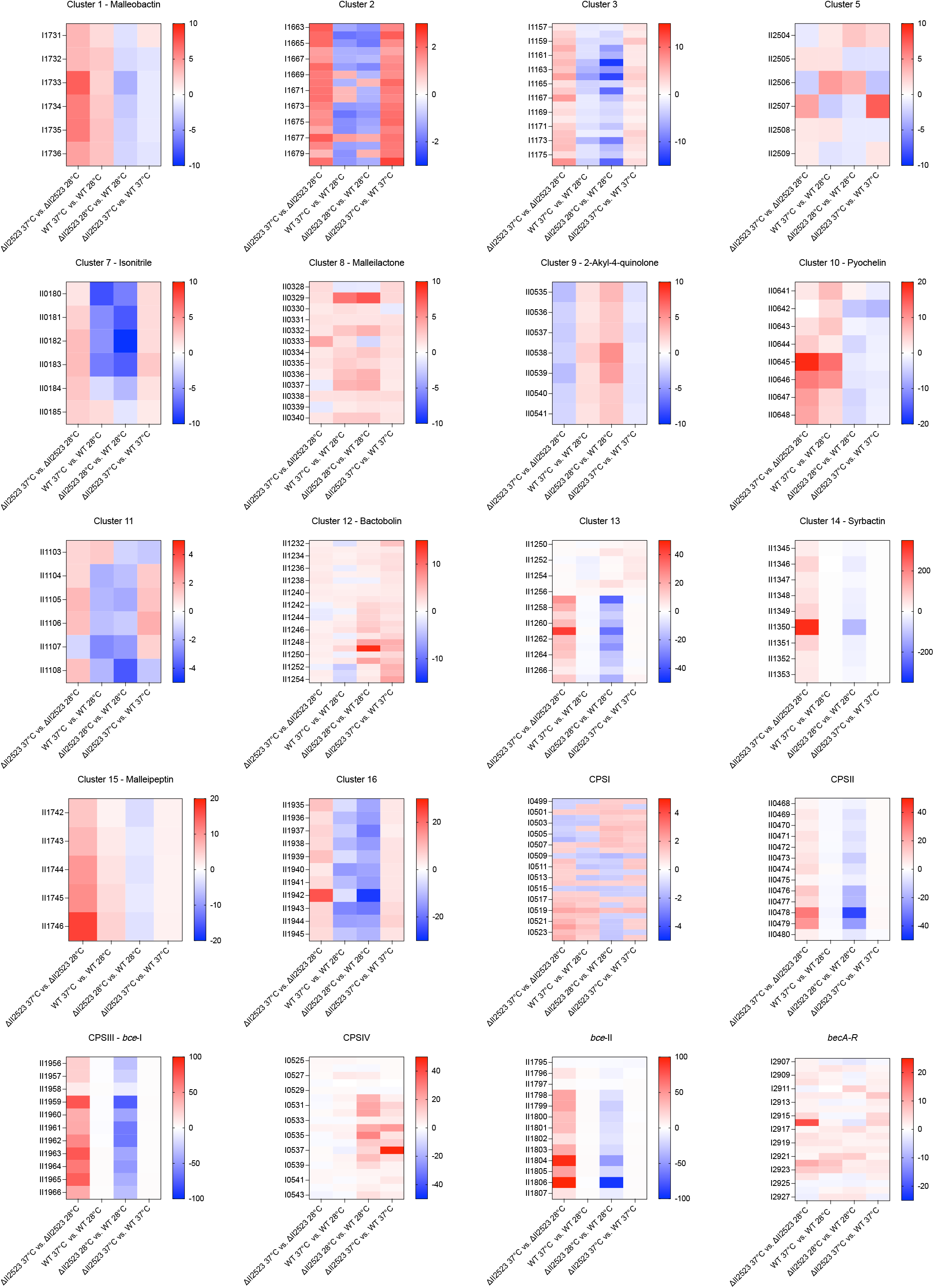
Heatmaps of complete predicted clusters show differential expression patterns of ΔII2523 in four pairwise comparisons. Raw fold change values for all predicted clusters described in this study were input from DESeq2. Color scales are consistent throughout (red being highest, white being neutral, and blue being lowest), although ranges vary to accommodate individual panels’ fold change differences.

**Fig S7.**
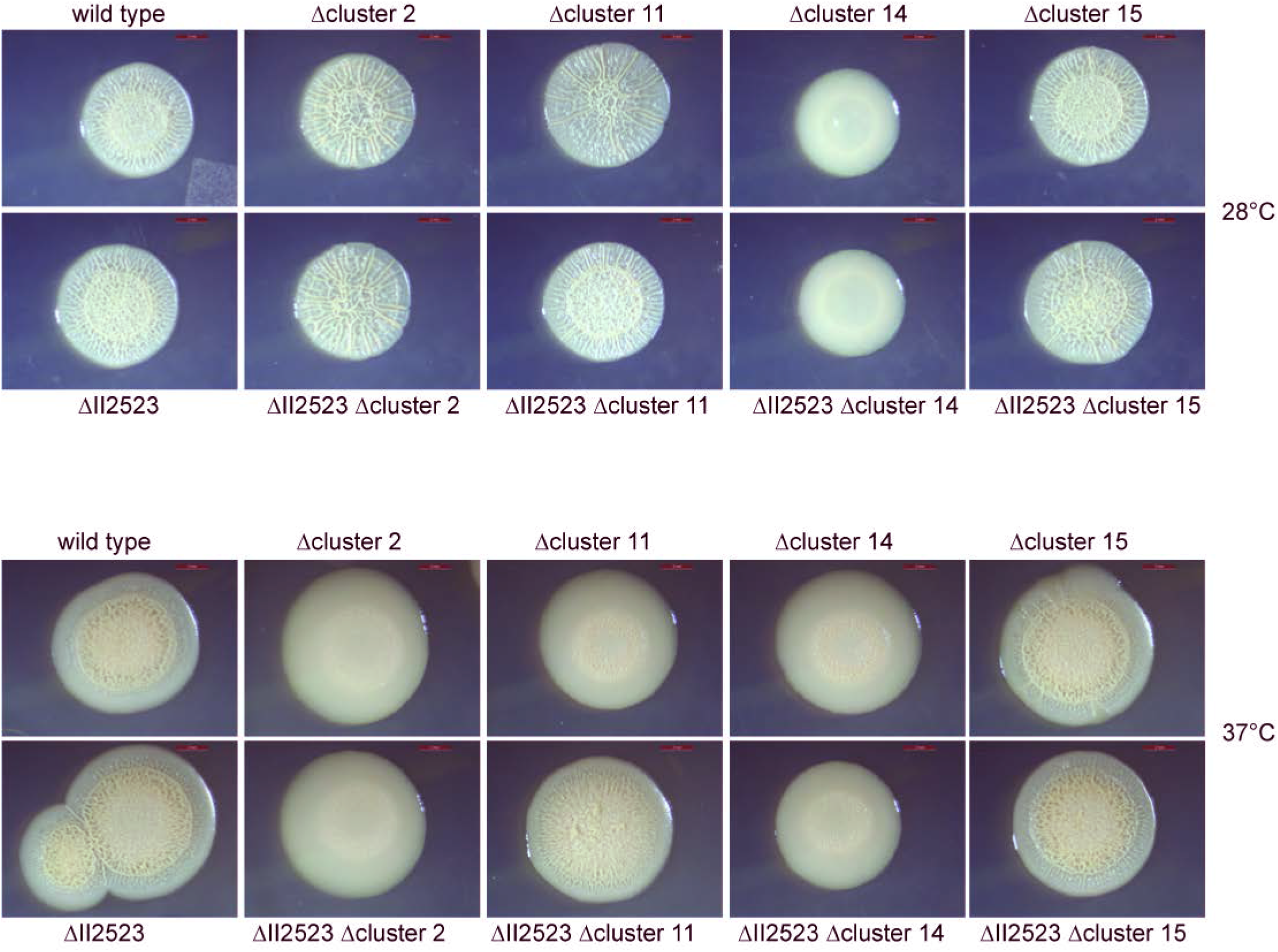
Colony morphology of NRPS/PKS mutants in Bp82 wild type or Bp82 ΔII2523 backgrounds on LB media at 28°C or 37°C. Images were taken after four days of growth. Scale bar represents 2mm.

